# Avalanches During Epithelial Tissue Growth; Uniform Growth and a Drosophila Eye Disc Model

**DOI:** 10.1101/2021.03.01.433318

**Authors:** George Courcoubetis, Chi Xu, Sergey Nuzhdin, Stephan Haas

## Abstract

In the physicists’ perspective, epithelial tissues constitute an exotic type of active matter with non-linear properties reminiscent of amorphous materials. In the context of a circular proliferating epithelium, modeled by the quasistatic vertex model, we identify novel discrete tissue scale rearrangements, i.e. cellular flow avalanches, which are a form of collective cell movement. During the avalanches, the cellular trajectories are radial in the periphery and form a vortex in the core. After the onset of these avalanches, the epithelial area grows discontinuously. The avalanches are found to be stochastic, and their strength is determined by the density of cells in the tissue. Overall, avalanches regularize the spatial tension distribution along tissue. Furthermore, the avalanche distribution is found to obey a power law, with an exponent consistent with sheer induced avalanches in amorphous materials. To decipher the role of avalanches in organ development, we simulate epithelial growth of the *Drosophila* eye disc during the third instar using a computational model, which includes both signaling and mechanistic signalling. During the third instar, the morphogenetic furrow (MF), a ∼10 cell wide wave of apical area constriction propagates through the epithelium, making it a system with interesting mechanical properties. These simulations are used to understand the details of the growth process, the effect of the MF on the growth dynamics on the tissue scale, and to make predictions. The avalanches are found to depend on the strength of the apical constriction of cells in the MF, with stronger apical constriction leading to less frequent and more pronounced avalanches. The results herein highlight the dependence of simulated tissue growth dynamics on relaxation timescales, and serve as a guide for *in vitro* experiments.

## Introduction

Ensembles of epithelial cells give rise to solid and liquid structural properties. Epithelial tissues are often considered viscoelastic materials, which exhibit elastic material properties on short time scales and liquid properties on long time scales (1). A prevailing notion is that the viscoelastic structure of tissues is a consequence of being in a liquid phase, but in proximity to a transition to a glass phase (2,3). Notably, growing and space adaptive tissues do not exhibit glassy behavior, since this would imply solid material properties on long time scales (2). Computational models that use Self Propelled Particle (SPP) models to describe epithelial dynamics capture a density dependent glass transition (2,4–6). However, such a transition can also be achieved by tuning cell structural parameters and fluctuations (7).

In this computational study, we find that the viscoelastic nature of tissue growth can be accompanied by sequences of cell flow avalanches. The tissue remains in an elastic solid structural state during growth. As the cells proliferate, the cell density increases until the internal pressure is sufficient to cause a large scale rearrangement event, which we define as an avalanche. During this rearrangement, the cells collectively flow radially, leading to a sharp increase in size. The properties of the identified avalanches are investigated in the context of a uniformly growing epithelial tissue and a model of *Drosophila* eye disc growth during the third instar.

In nature, avalanches are ubiquitous. From earthquakes to ferromagnets, large scale structural rearrangements arise, whereby a small perturbation leads to a pronounced collective response (8,9). Systems characterized by such avalanche instabilities exhibit self organized criticality (SOC) (10). A main result is that SOC systems give rise to power law scaling signatures in observables, and by fitting to power laws, universal exponents can be obtained. These inferred exponents can be used to identify the strength of correlations in the system and the universality class of the process. Universality classes are groups of processes that share the same scaling properties. As an example, amorphous materials, which include glassy systems, exhibit avalanche behavior when exposed to strain (11,12). A successful characterization of such avalanches utilizes the physics of self organized criticality and phase transitions (11,13). Recently, sheer stress induced avalanches were identified in the context of a vertex model, and both their universality class and material properties were mapped to amorphous materials (14). Furthermore, signatures of criticality were identified *in vitro* for the *Drosophila* wing disc epithelium, suggesting that the nonlinear properties of collective processes in amorphous materials and the vertex model are at play in biological tissue growth (14).

Epithelial tissue growth and migration has been the subject of several recent *in vitro* studies. These generally branch into two categories, either focused on confined tissues (15–18), or on freely expanding tissues (19–22). In the former category, processes that involve density induced motility transitions, front propagation, and spatial sensing are probed. In the latter category, processes involving mesenchymal-to-epithelial transition, contact inhibition, size dependence of growth and boundary formation are investigated. In both categories, cell motility and collective phenomena are a central theme. In the context of growth, a ubiquitous collective phenomenon, identified *in vitro* for both freely expanding and constrained growing cell tissues, is the formation of vortices by cell trajectories (18,20). Similar to our discovery of avalanches during growth, which is reported here, the vortices are a density driven collective phenomenon (18). Interestingly, the cell trajectories during avalanches also exhibit vortex signatures. However, the avalanches also involve a coordinated radial motion of cells not reported in (18,20). In our study, the underlying model treats the tissue in a confluent state, as in epithelial tissues *in vivo*, where growth occurs in compartments separated by boundaries. Thus, it provides an appropriate framework linking the phenomena probed with what takes place *in vivo*.

Biophysical properties, such as mechanical forces, impact cellular processes during growth phases in development (23). Epithelial cells have been shown to regulate their shape and division rate by responding to mechanical force stimuli (15,24–26). Furthermore, control of the cell division plane via cell-shape and mechanical considerations has been experimentally detected in animal and plant epithelial cells (24,27). Numerical implementations of epithelial tissue growth, which include mechanosensing and division plane control, have shown to generate improved epithelial mosaics that represent *in vitro* structures more closely (28,29). Single cell resolution computational models are a valuable tool to model growth, since each cell can respond to its distinct state and incorporate phenomena at the single cell level (30,31).

We use a vertex model (32) to simulate the spatial dynamics of epithelial tissues. Vertex models have provided successful mechanical simulations of confluent epithelial tissues with single cell resolution (33,34). This type of epithelial tissue computational modeling treats cells as polygons and uses energy considerations to track the spatial arrangement of cells, vertices, and edges. It has been successful in simulating apoptosis, mitosis, and cell shape changes (33,34). In the vertex models, each edge, perimeter and area of the cells contribute to the energy and affect the structure of the tissue. There are two main categories of dynamics implemented in vertex models. The first kind is the active vertex model, which explicitly updates the structure using the equations of motion for each cell or vertex in the overdamped regime (35). The other kind is the quasistatic vertex model, which assumes that relaxation occurs in very short time scales and drives the system to the first accessible local energy minimum using gradient descent (32). The local energy minima correspond to mechanically stable configurations, as the negative of the gradient of the energy corresponds to the forces on the vertices. These tissue models have been verified in Drosophila development with experimental data and laser aberration experiments, and have been recently successfully implemented in describing complex phenomena such as epithelial tissue folding and growth (32,35,36).

Epithelial tissues, across metazoa, have characteristic cellular packing geometries and distributions captured by the vertex model. One characteristic of epithelial geometry is formulated in Lewis’ law, which states that the average cell size for cells with n neighbors, *Ā* _*n*_, is proportional to the number of neighbors (37). This characteristic distribution has been found in a variety of metazoan tissues (32,38–40). Another characteristic signature prevalent across growing epithelial tissues is the distribution of cell sides (28,29,32,41). The cells are predominantly hexagonal, with pentagons and heptagons following suit. These experimentally observed cellular distributions have been reproduced by models that combine single cell geometrical configurations and proliferation (28,30,32).

To investigate epithelial tissue growth in the context of an *in vitro* system, we simulated epithelial growth in the *Drosophila* eye disc during the third instar. The *Drosophila* eye disc is part of the eye-antennal disc, which begins with approximately 70 founder cells and develops to a final size of approximately 44,000 cells (42). During the first half of the second instar, the eye-antennal disc grows via rapid cell proliferation, and at the late second instar, it segregates into the retinal progenitor and the antennal/head epidermal fields (43). Photoreceptor development initiates at the beginning of the third instar, on an elliptical epithelial sheet, the retinal progenitor, composed of undifferentiated cells. The size of the retinal progenitor starts at approximately 10, 000 μ*m*^2^ (44). At that stage, a visible indentation of the tissue, termed the morphogenetic furrow (MF), begins propagating linearly from the posterior to the anterior of the eye disc, leaving behind differentiating photoreceptors (43,45–47). Cells within the MF region are constricted, with their apical surface reduced by a factor of about 15, from 9 μ*m*^2^ ahead of the furrow to 0.6 μ*m*^2^ (47). There are two mitotic waves, the first mitotic wave occurs anterior to the MF, where cells differentiate asynchronously, and the second mitotic wave affects undifferentiated cells between photoreceptor cells behind the MF (47,48). The first mitotic wave produces the majority of cells of the final eye disc and takes place ahead of the MF, whereas the second mitotic wave ensures that there are sufficient cells for proper ommatidial formation (42). The second mitotic wave is more controlled, takes place posterior to the MF, and occurs in a synchronized and organized manner (49). The contribution to growth of the second mitotic is negligible compared to the first mitotic wave and will not be included in the model (44).

The growth dynamics of the third instar *Drosophila* eye disc have been described quantitatively. The growth rate over time has been determined by analyzing experimental tissue size data over different developmental time points (44). Both exponential decay and inverse area functional forms fitted the data exceptionally well (44). Furthermore, the area rule has been the focus of a subsequent study, which identifies a diluting and decaying transcript, cytokine unpaired, as a possible candidate for cell growth regulation (50). Thus, functional forms and coefficients that describe the time dependent eye imaginal disc growth rate quantitatively are available and were used to simulate the process.

In order to simulate the propagation of the MF, cellular signaling was incorporated on top of the vertex model. *In vitro*, the MF is a result of signal transduction of multiple morphogens with multiple roles. The primary signal that is associated as the initiator and driver of the MF is hedgehog (Hh) (43,46,51–53), a morphogen that is sourced from differentiating neural cells posterior to the MF. Hh diffuses in the epithelial tissue up to and including the constricted cells in the morphogenetic furrow (46). The propagation of the MF is mediated by Decapentaplegic and regulated by Hemothorax, Hairy and extra macrochaetae (46,54–56). In the model herein, a simplified gene regulatory network was used, see (57) for a model with a detailed treatment of the gene regulatory network. The propagation of the MF was simulated by introducing an effective morphogen that induces its own expression while diffusing from posterior to anterior (see methods for details).

Few studies have so far been published that combine signal transduction processes, transcriptional responses and vertex models (31,58–61). In the model by Wartlick et al, the authors were able to reconstruct the observed growth of the wing disc via an integrated signal transduction and the vertex model by Farhadifar et al (61). Schilling et al. developed a model for the production, diffusion, and sensing of Hh in the Drosophila wing primordium and explained how anterior/posterior cell sorting can be explained by a homotypic boundary model (59). We will use a similar approach, i.e. a finite-volume method defined by a tessellation of the centroids of the cells. Notably, an alternative approach has been introduced utilizing the Lagrangian–Eulerian (ALE) formulation and finite element method to describe morphogen transport within a tissue coupled to a vertex model (60). Furthermore, an earlier study was able to model wing disk growth by considering a complex mechanical feedback, with cell constriction affecting the cell cycle and a biased angle of mitosis (30). Finally, a report on the zebrafish retina was able to model highly ordered packing via coupling planar cell polarity proteins with edge tension (31,33). The eye disc model developed here belongs in the class of models that couple signaling and vertex models. In the context of the eye disc, the model is novel in its treatment of the epithelial structure and a natural extension of the viscous fluid based treatment of drosophila eye disc growth seen in (57).

## Results

### Generic uniform growth model

In order to study cellular avalanches during epithelial tissue growth, a vertex model is used, which is based on an energy functional that optimizes cell surface area, and minimizes edge tension as well as contractility of the cell perimeter. In our initial implementation, the tissue is initialized to a small circular size and is allowed to proliferate into empty space. The cell doubling rate is set to be the same for each cell and constant over time. In addition, the doubling rate is also set to weakly depend on cell area (29), approximating cellular mechanosensing, with bigger than average cells having a higher division rate compared to smaller cells (see methods for details). Contrary to the periodic boundary condition case, we discovered the addition of size dependent doubling rate was necessary for open boundary growth (SI Fig. 1). Without mechanosensing, the cells become unrealistically packed, there is significant apoptosis and the increase of tissue size is suppressed. Importantly, avalanches were detected with or without the size dependent doubling rate. Furthermore, the tension at the tissue peripherie is set to an increased value in order to enforce a smooth boundary. The approximately circular symmetry of this system allows for probing the radial profile of structural quantities. Three frames at different times of a simulation realization are shown in Figs. 1A-C. In Fig. 1C, the tissue increases significantly in size, as a constant cell division rate leads to exponential growth. A complete video of a simulation of this growth process can be found in SI Vid 1.

**Fig. 1.**
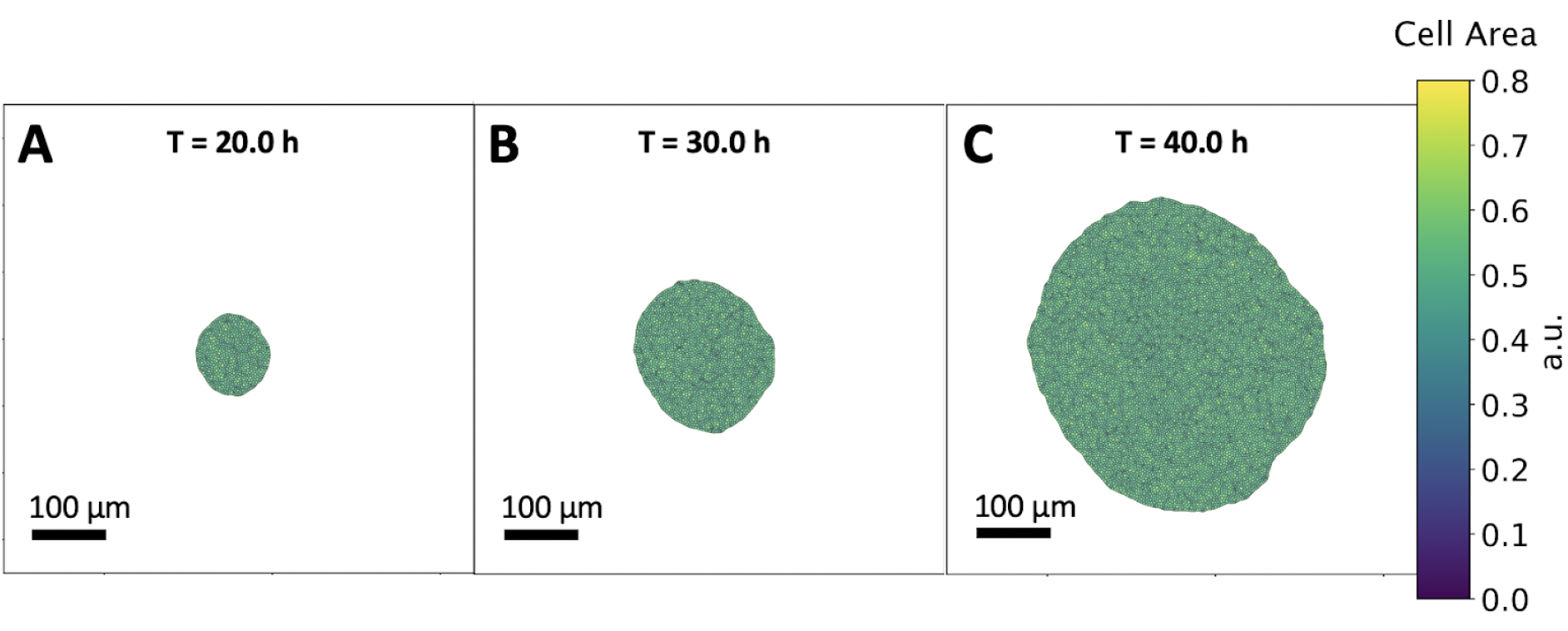
Uniform growth model. **A-C:** Successive frames of the tissue growth process at different time points, T=20.0h, 30.0h and 40.0h respectively. The tissue starts at a small size composed of 50 cells and is simulated until it reaches 350 times its original size, allowing the understanding of different growth regimes. Cell division takes place at a constant doubling rate of ∼6hrs. The cells are color coded with respect to their area. The tissue retains a smooth boundary during growth and an approximately circular shape. In early times, the tissue grows continuously and exhibits discontinuities, or avalanches, at later times of ∼40 hrs.

Next, we probe aspects of the collective cell motility during a characteristic avalanche. In Fig. 2A, the cell trajectories during an avalanche event are visualized. During the avalanche, the cells move collectively, with displacement increasing in a radial fashion. Interestingly, the trajectories of the cells also have a significant tangential component. The tangential component is visualized in Fig. 2B by color coding the trajectories with their respective angles. Trajectories of cells in the middle of the tissue have a significant tangential component, contrary to the radial direction of trajectories on the periphery. This signature indicates the presence of a cellular vortex. Interestingly, tissue scale vortex formation has been experimentally observed in growing epithelial monolayers (20). In addition, the radial profile of the vortex, formed during an avalanche, is consistent with the profile reported experimentally (20). For the cell trajectories without the onset of avalanches, see SI Fig. 2.

**Fig. 2.**
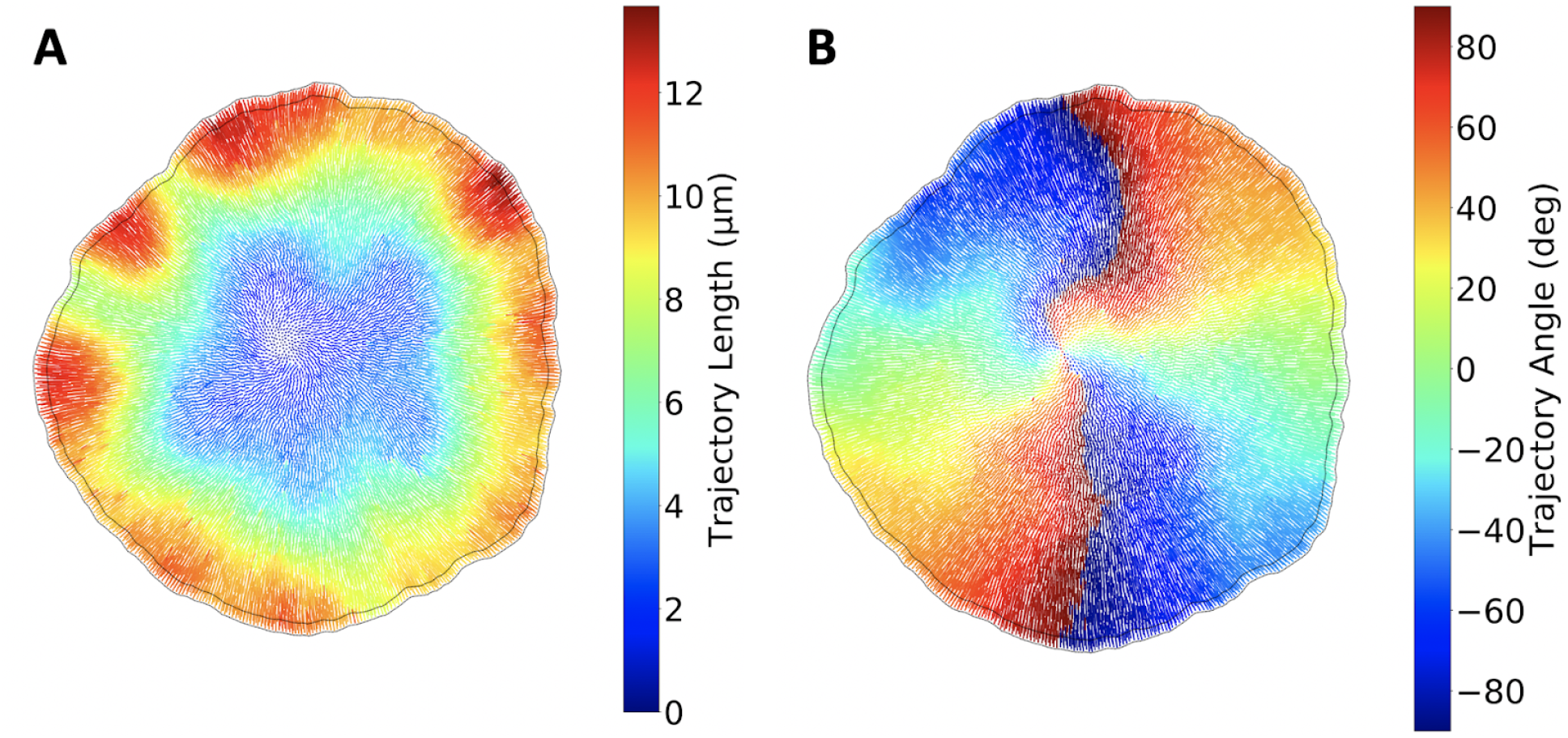
Single cell trajectory characteristics during avalanches. **A:** Cell trajectories during an avalanche, color coded with their respective length. The cells in the outer region displace the most, whereas the trajectories are shorter at the center of the tissue. The tissue boundary before and after the avalanche is outlined, visualizing the net expansion of the tissue. **B:** Cell trajectories during an avalanche, color coded with their respective angle relative to the x-axis, as defined by the inverse tangent function. On the outer region of the tissue, the trajectories are approximately radial. The vorticity is evident towards the center of the tissue, where the trajectory angles start to change.

The time dependence of the evolving total area and average cell area in this realization is depicted in Figs. 3A&B. Initially, the tissue grows at a predominantly continuous rate, as cells can easily rearrange within the tissue. As it becomes larger, at approximately 30h, more pronounced discontinuities in the area growth are detected. These avalanches correspond to large scale cell rearrangements. Due to the initial small size of the tissue, the increased boundary tension dominates the energy term, and the tissue begins its growth at a high cell packing state. As the tissue grows further, the average cell area increases, as the boundary tension energy becomes negligible compared to the bulk energy.

**Fig. 3.**
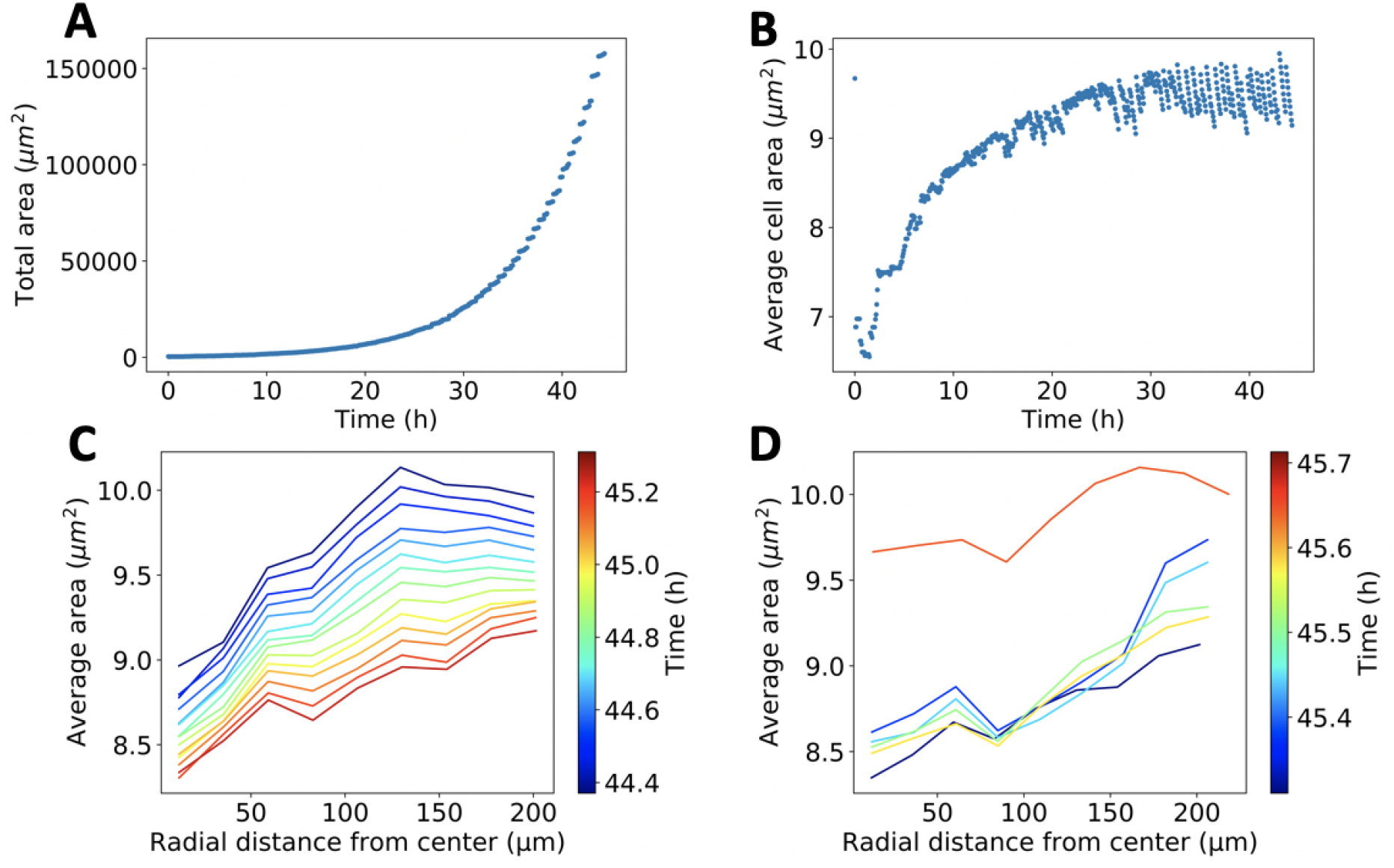
Uniform growth model observables. **A:** Total tissue area versus time. The tissue is initiated as a small size circle, and the cells undergo mitosis with a constant doubling rate. At early times, the tissue follows predominately continuous growth. After ∼30 h, the tissue displays growth predominately with discontinuities. **B:** Average cell area versus time. The discontinuities in average cell area become more pronounced and regular at ∼30 h. **C-D:** Avalanches in the uniform growth model. **C:** Average cell area versus radial distance from the tissue center at consecutive time points. The spatial signature of the average cell area before an avalanche occurs is depicted. As cells divide, the average cell area decreases globally, with higher packing from the center to the tissue boundaries. **D:** Average cell area versus radial distance from the tissue center at consecutive time points. The spatial signature of the average cell area is depicted shortly before and after an avalanche occurs. Before the avalanche, there is a positive gradient from the tissue center to the tissue boundaries for cell size. After the avalanche, the average area increases with a discontinuity. The magnitude of the gradient diminishes.

The radial profile of the average cell area is probed to understand the onset of avalanches. Before the avalanche, the cell density increases as cell division takes place (Fig. 3C). The spatial signature suggests increased cell packing in the center of the tissue, which diminishes outwards. Right after the avalanche, the radial cell density discontinuously increases (Fig. 3D), and the gradient of cell packing is less pronounced after the avalanche. This is consistent with the radial increase of trajectory length shown in Fig. 2A.

Based on these measured quantities, we can address what causes the avalanches and what controls their magnitude. To this end, we repeat 150 simulations of the process and isolate the avalanches. To do so, we skip the first 30 h of development and record an avalanche if a cell density discontinuity is equal to 0.1 μ*m*^2^ or greater. To quantify the spatial inhomogeneity of cell packing, we use the standard deviation of the average area as a function of radial distance from the tissue center. Interestingly, Fig. 4A shows that there is a strong correlation between the average cell area before the avalanche and the jump magnitude, with a spearman’s rank correlation coefficient of ρ=-0.83 and a p-value of 0. Furthermore, SI Fig. 3 illustrates that there is weak correlation between spatial density inhomogeneity and jump magnitude. Thus, avalanche onset and strength are more sensitive to the global density of cells rather than to the details of their spatial distribution.

**Fig. 4.**
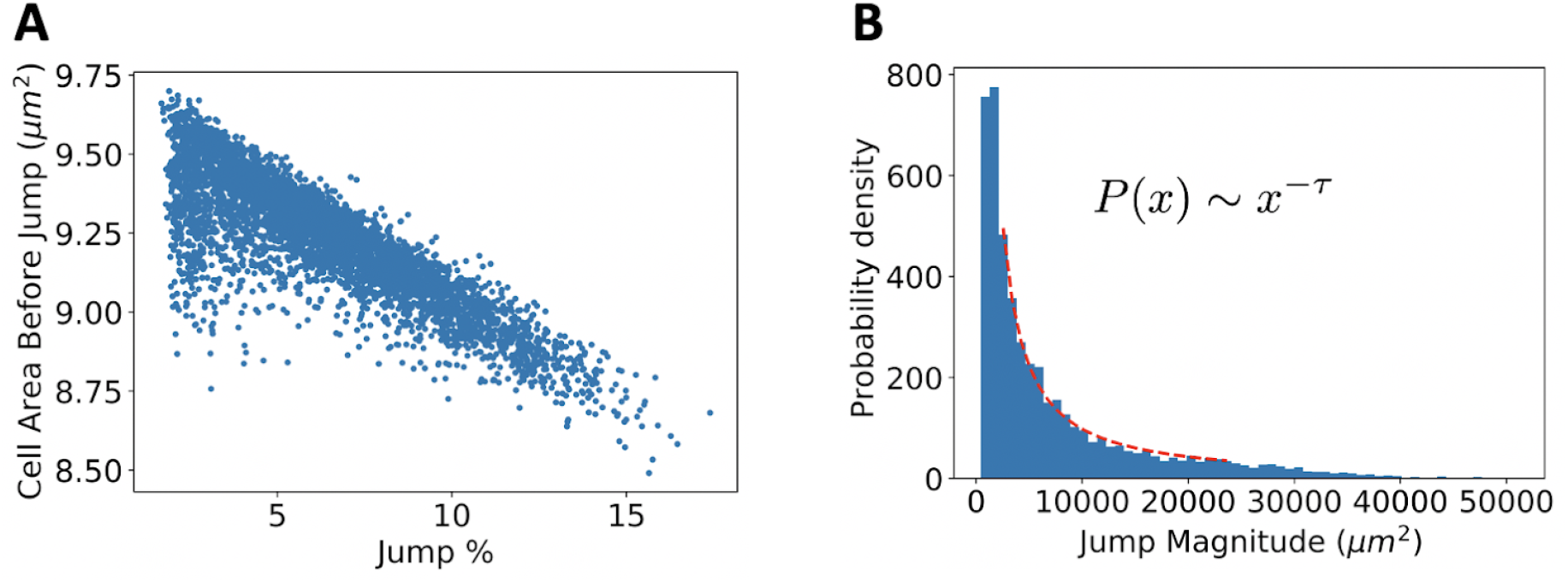
Avalanche Characteristics. **A:** Average cell area versus avalanche magnitude. The avalanche magnitude is quantified as the percent discontinuity in the total area. The more tightly packed the epithelium, right before the onset of the avalanche, the larger the total area discontinuity. **B:** Probability density of avalanche magnitudes. The functional form is a power law distribution, with a fit shown by the red dotted line. The exponent found is τ=1.27 ± 0.04. Such power law distributions are a universal characteristic of avalanches found in nature.

To understand whether the system exhibits self organized criticality (SOC), the probability distribution of avalanche magnitudes is shown in Fig. 4B. To provide more avalanches and solidify the fit, the growth model was run 150 times for an extended runtime, until the tissues reached 300, 00 μ*m*^2^. The magnitudes are distributed according to a power law distribution, with an exponential cutoff, which is a ubiquitous property of SOC. The scaling exponent is found to be τ=1.27, in agreement with analogous exponents observed in sheer induced, stress avalanches in overdamped elastoplastic models (11,62). More information about the fitting procedure and comparison with exponential fit is shown in SI Fig. 4. This result reinforces the notion that the vertex model and epithelial tissues share common material properties and universality classes with plastic amorphous systems (14). Varying the boundary tension and the mechanosensing amplitude did not cause the exponent τ to change (SI Fig. 5).

**Fig. 5.**
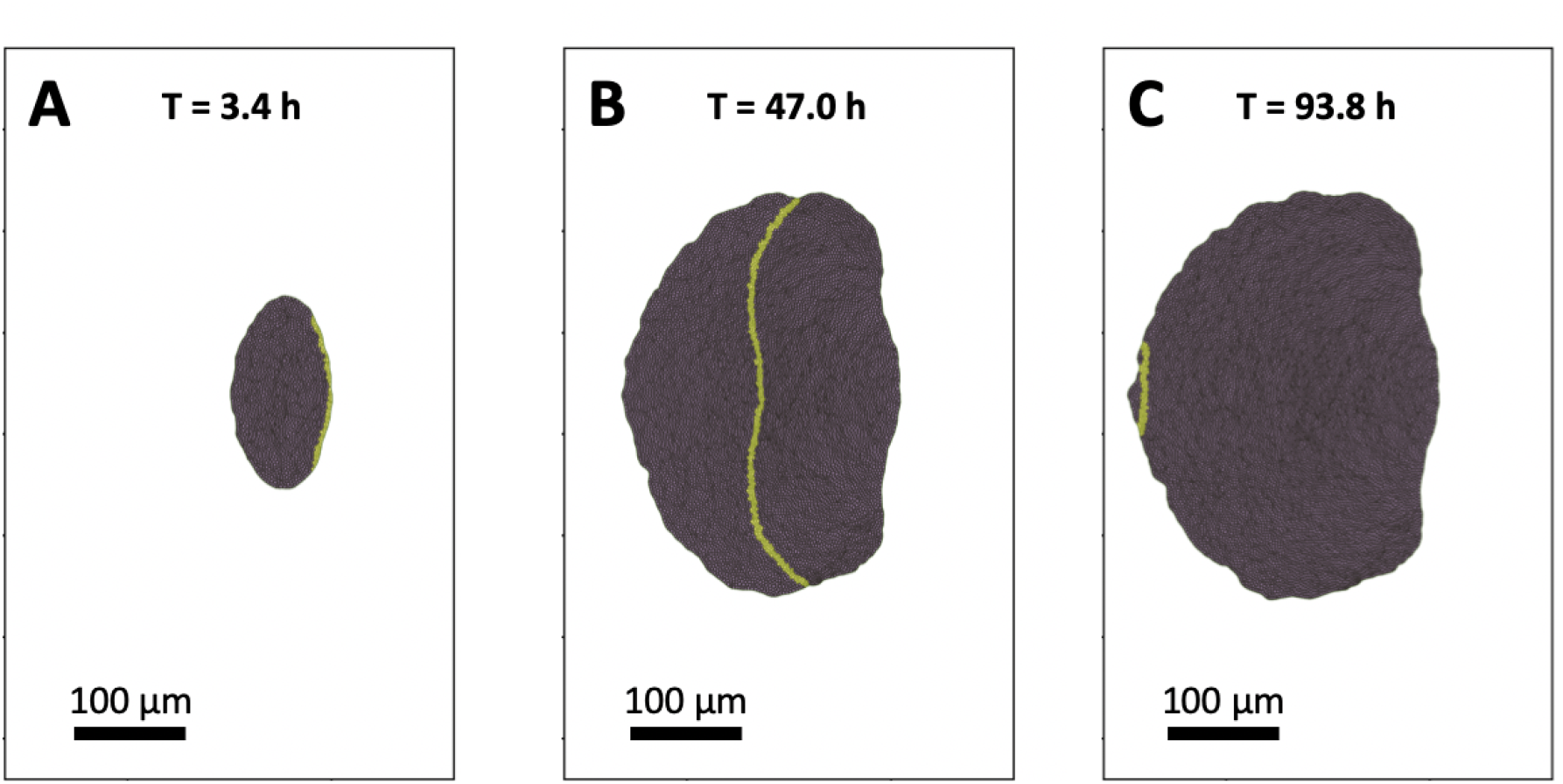
Drosophila eye disk tissue at different times of growth. **A-C:** Vertex model simulation of tissue growth, with cells in the MF highlighted in yellow. **A:** Time T=3.4 h; the MF is initialized at the posterior tissue boundary. **B:** Time T=47.0 h; the majority of cell divisions have taken place, while the MF has linearly propagated to the bulk of the tissue. **C:** The MF reaches the anterior boundary of the tissue, signaling the cell cycle arrest of all cells and termination of growth.

### *Drosophila* eye disc growth model

In order to study the cellular avalanches in the context of a particular developmental process that is experimentally accessible, a more detailed model was developed. In this version of the model, the cell doubling rate is set by experimental data, the tissue is initialized in an elliptical shape, and there is signaling of a single effective morphogen which drives the propagation of the morphogenetic furrow (MF) see methods for details. The boundary tension at the boundary is again set to the same increased value as the original uniform growth model to ensure a smooth boundary. A complete video of a simulation of this growth process can be found in SI Vid 2.

A visualization of the simulation of the growth process at three different time steps, initiation, middle, and end of growth are shown in Fig. 5. Initially, the transcription factor is induced in the posterior boundary cells (Fig. 5A). This will eventually cause the auto-activation of transcript production in neighboring cells, and will lead to the initiation of the MF. At this time point, all the cells are dividing in the tissue. At the midpoint, the MF has traversed approximately halfway through the tissue (Fig. 5B). Cells entering the MF immediately stop dividing and become immobile 2.5 h after the MF has passed, to simulate differentiation. At the midpoint, the growth rate has decreased by approximately 80%, and the majority of cell divisions has taken place. Finally, at the end point (Fig. 5C), the growth process is complete, as the MF has propagated to the anterior tissue boundary, and all cells have arrested their cell cycle.

During the growth process, the structural details of the cellular mosaic of the tissue were measured. To quantitatively describe the mosaic of the epithelial cells, average areas versus number of sides and the distribution of polygon classes were plotted (Fig. 6). The measured quantities match experimentally established, system independent, epithelial structure signatures. The emergent relationship between average cell size of a given polygon class, versus number of edges is linear (Fig. 6A), in accordance with Lewis’s law. Furthermore, the distribution of cell polygon classes matches the corresponding distribution found *in vitro* (Fig. 6B) with P=0.98. Thus, the model gives rise to realistic epithelial packings.

**Fig. 6.**
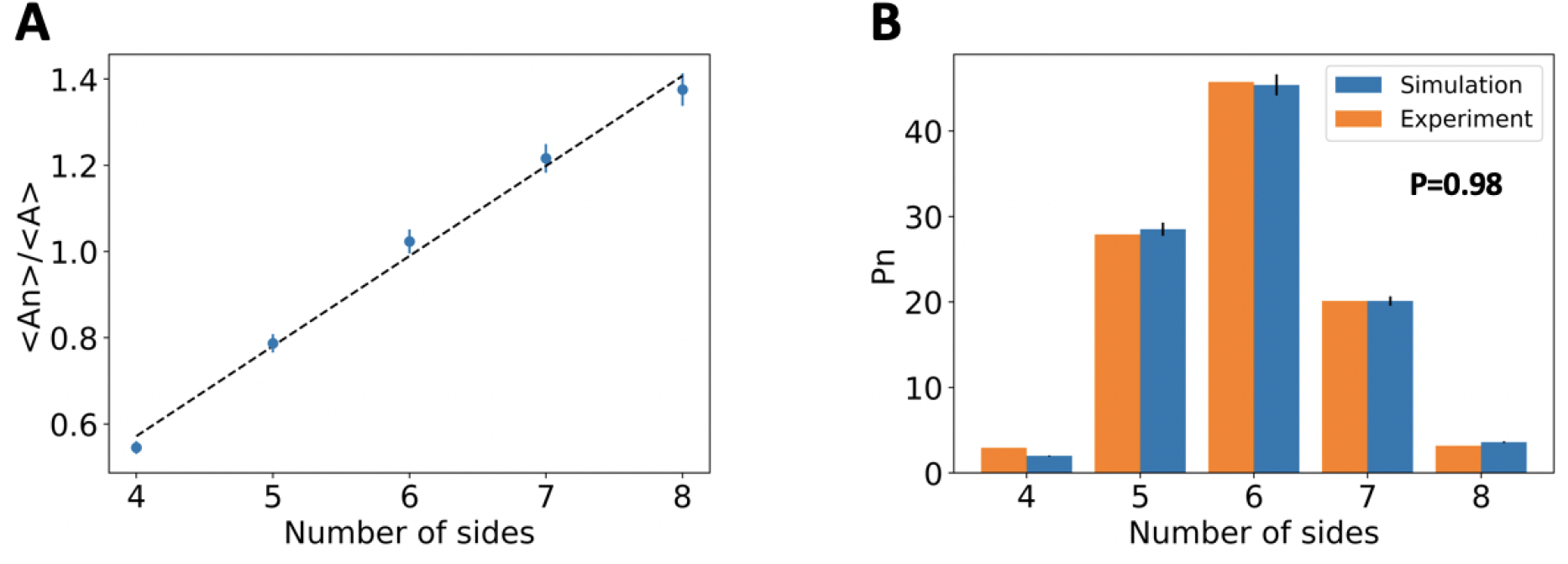
Quantifying epithelial packing during the simulated growth process. **A:** Average cell area per number of cell sides, divided by the mean area, is plotted versus cell number of sides. The values are obtained by combining data from all frames in the simulation. A linear relationship is observed, in accordance with Lewis’ law. **B:** Distribution of number of cell sides. The simulation values are obtained by combining data from all frames in the eye disc simulation. The experimental values were obtained from (41) and correspond to *Drosophila melanogaster* third instar larval wing disc.

Aspects of the tissue growth process can be captured by quantifying the temporal evolution of key observables. The anterior area depicted in Fig. 7A initially grows, reaches a plateau and then decreases until it vanishes, as needed for growth termination. There are three effects contributing to the anterior area profile. Firstly, new cells are introduced due to cell mitosis. Second, since the MF separates the anterior from the posterior compartment, as the MF moves across the tissue, it turns anterior cells to posterior cells. Third, the rate of mitosis in the anterior decreases over time. The posterior area depicted in Fig. 7C grows solely via the cells that exit the anterior area due to MF propagation. Initially, the eye disc is small, and the MF passes through a small number of cells. At the midpoint, the posterior area grows the fastest, as the MF is at the center of the tissue and passes through a large number of cells. Towards the end of the growth process, it tapers down, as it reaches the anterior edge. The total area depicted in Fig. 7B is the sum of the anterior and posterior compartments. Finally, the posterior length depicted in Fig. 7D, is shown to increase linearly over time. It is defined as the length from the posterior edge midpoint to the midpoint of the MF. Initially, there is a plateau, which corresponds to the initialization time needed for the MF to begin propagating. Casting aside the initialization time, the linear nature of the posterior length (*R*^2^ = 0.99) indicates that the posterior length indicates that the MF is propagating at a constant speed throughout the growth process. The value of the slope of the posterior length corresponds to the MF speed, which is 3.4 μ*m/hr*, matching the experimental value.

**Fig. 7.**
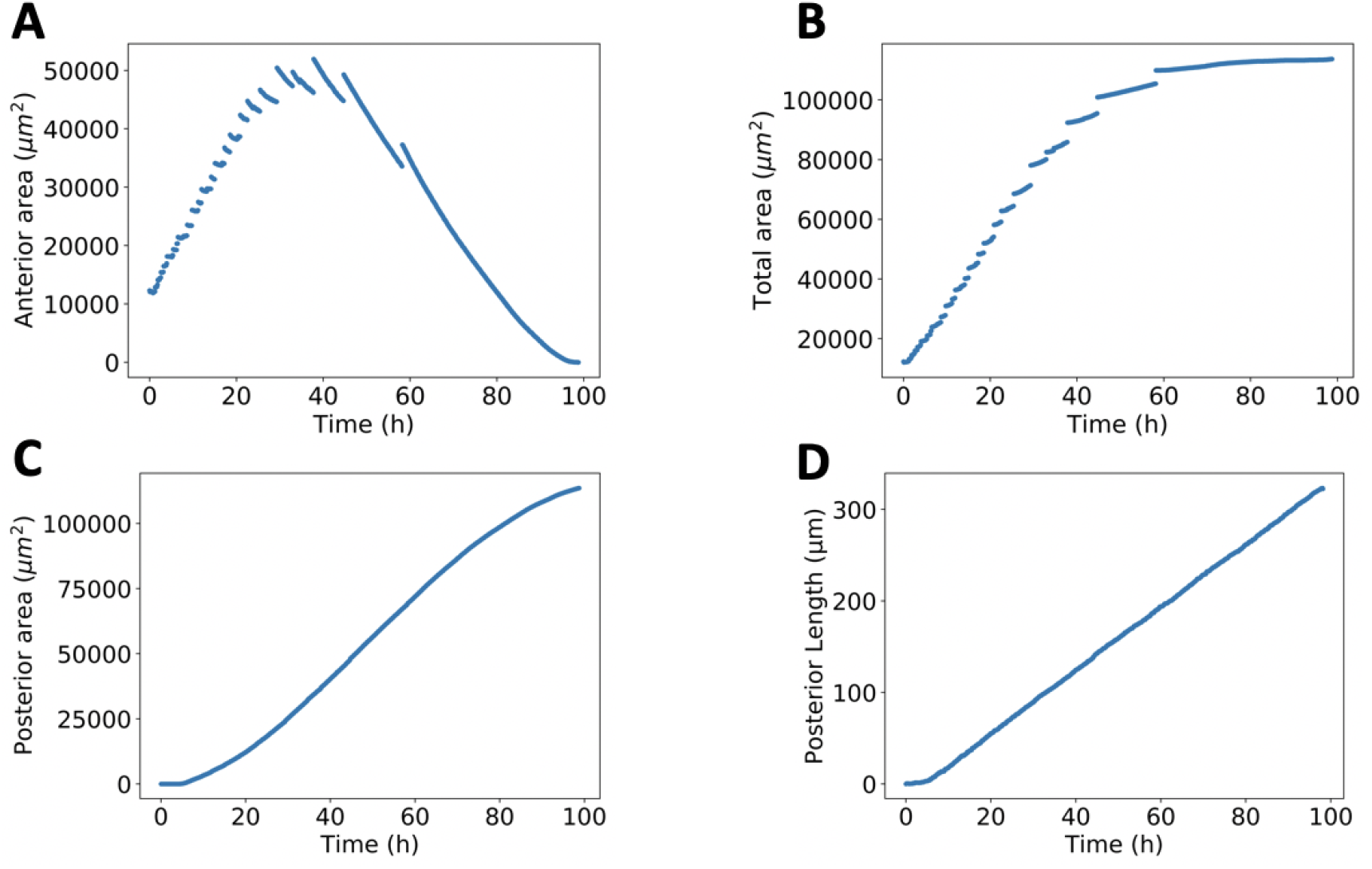
Drosophila eye disc growth dynamics. **A:** Time dependence of anterior area, defined as the portion of the tissue ahead of the MF. Cell division takes place only on the anterior portion of the eye disc. **B:** Time dependence of total area. The final value is in the same order of magnitude found in wild-type *Drosophila* strains. **C:** Time dependence of posterior area, defined as the portion of the tissue before the MF. Cell division is arrested in the posterior portion of the tissue. The posterior grows solely by cells exiting the propagating MF. **D:** Time dependence of the posterior length, defined as the distance of the posterior edge to the midpoint of the MF. Aside from initialization time, the posterior length increases linearly, with slope of 3.4μm/hr, in accordance with MF propagation experimental data.

Similarly to the simpler model, the quantification of the growth process reveals discontinuities, i.e., avalanches, in the temporal evolution of the tissue area. This can be seen in the anterior area in Fig. 7A, and consequently also in the total area (Fig. 7B). To understand better what is taking place at the single cell level and across tissue compartments, the average cell area during the growth process is plotted in Fig. 8A. In Fig. 8B we visualize the tissue area avalanche magnitude for each tissue compartment. Comparing Fig. 8A to Fig. 8B, one can see that each avalanche is accompanied by a discontinuity in the average cell area. Furthermore, the discontinuities take place after the average cell area decreases and always lead to an increase of the average cell area. Therefore, the avalanches are a consequence of increased cell packing in the eye disc that leads to a global rearrangement of cells. Also, the avalanches mainly take place in the anterior of the eye disc (Fig. 8B).

**Fig. 8.**
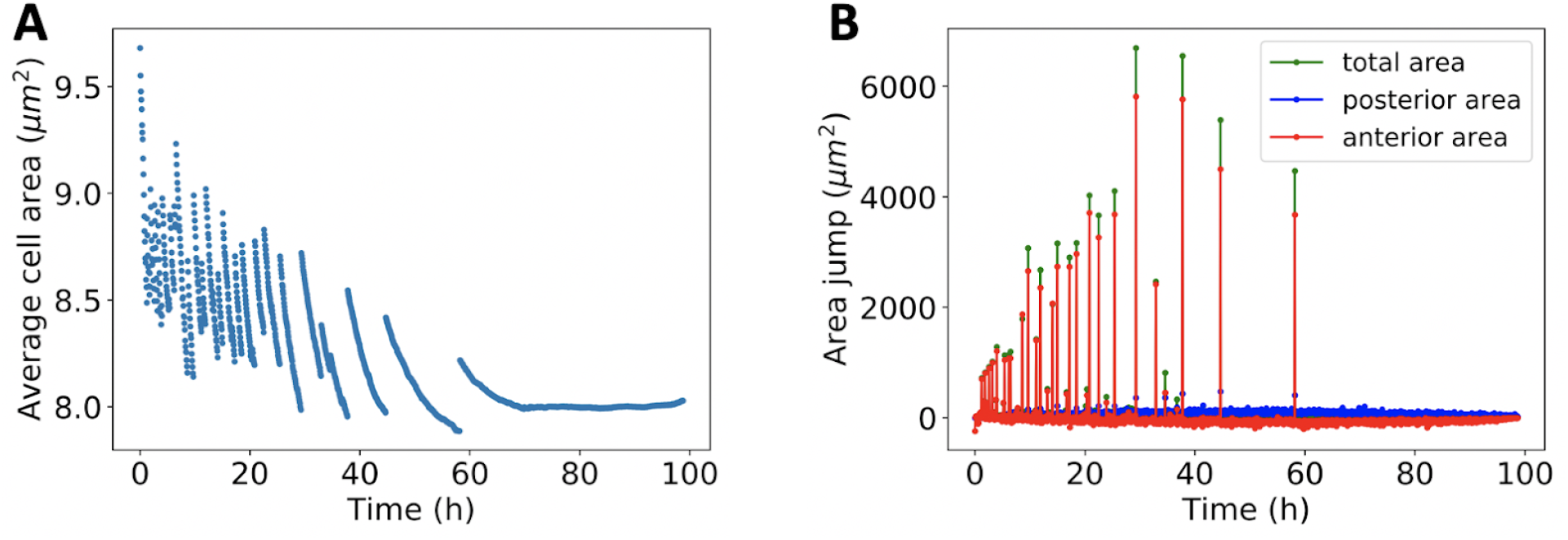
Avalanches in eye disc growth dynamics. **A:** Time dependence of the average cell area. The avalanches are captured by discontinuous transitions from high cell packing to low. **B:** Time dependence of tissue area jumps for total, anterior and posterior area, marked by green, red and blue respectively. The avalanches lead to tissue area discontinuities, with the majority occurring in the anterior.

Changing apical constriction affects the onset of the avalanches. In the model, we can tune the strength of apical constriction in the MF. In Fig. 9, the MF cell constriction is incrementally increased from no constriction to high constriction, by changing the preferred area of cells in the MF. We define the apical constriction factor as the percent decrease in the preferred cell area of cells in the MF. Fig. 9A shows the effect of the constriction strength on the cell number, with higher constriction leading to significantly larger cell numbers. On the other hand, Fig. 9B shows total area, which reflects that eye discs are larger when the apical constriction in the MF is increased. However their relationship is not a scalar multiple, as average cell size is variable between different apical constriction strengths. This can be seen by inspection of two simulation examples (Figs. 9C & 9D) of the average cell size, one for no apical constriction and one for strong apical constriction in the MF. When the apical constriction is increased, the avalanches become less frequent and the average cell area before they take place is decreased. Thus, apical constriction makes it harder, in the sense that cells reach a higher packing state, for global cell rearrangements take place. Finally, since the reaction-diffusion equation is discretized on the lattice of the cells, independent of packing, a decreased cell size leads to a decrease in the MF speed (SI Fig. 6). Since the growth rate decays based on the posterior length and thus the MF speed, apical constriction makes the growth rate decay slower.

**Fig. 9.**
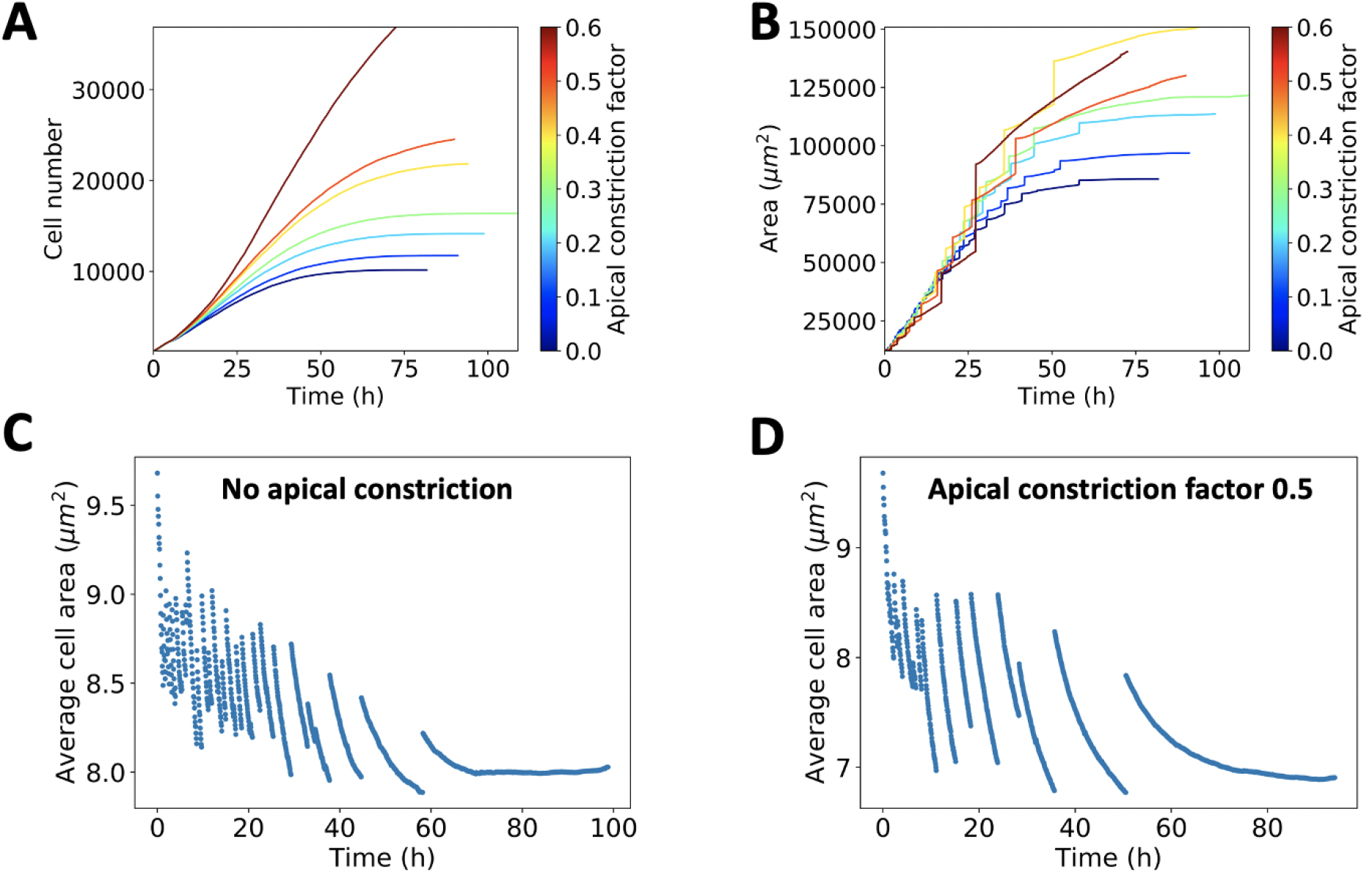
Avalanches depend on the magnitude of apical constriction in the MF. **A:** Time dependence of the cell number. The larger the apical constriction, the higher the cell number. **B:** Time dependence of the tissue area for different constriction strengths. The larger the apical constriction, the bigger the tissue area. **C-D:** Average area for no constriction and moderate constriction. The case of no apical constriction corresponds to more frequent avalanches that take place in higher densities. This highlights the observation that apical constriction in the MF makes avalanches harder to achieve, and leads to higher cell packing.

### Model Predictions

The model predicts that proliferating epithelial tissues, with short relaxation time scale dynamics, exhibit discontinuous spurs of growth. The onset and strength of the discontinuities, which we term avalanches, are positively correlated with increased cellular packing. Thus, the model predicts that the average cellular area depends strongly on whether the tissue growth is before or after an avalanche. Furthermore, the model predicts the formation of tissue scale vortices during growth, in agreement with experiments (20).

In the context of the *Drosophila* eye disc development, with the assumption that the dynamics fall into a regime of short relaxation time scales, we expect that the average cell density will exhibit considerable variation during growth. Since avalanches take place less frequently at later times during tissue growth, the variations should be less intense at later developmental times, consistent with a more densely packed eye disc. In addition, the stochastic onset and strength of avalanches suggests that cellular density should vary between eye discs at the completion of eye disc growth. Increase of the apical constriction in the MF makes avalanches harder to attain, with cells needing to be more packed. Thus, mutant eye discs with lower apical constriction should, on average, exhibit a larger average anterior cell size compared to the wild-type.

## Discussion

To date, most insights regarding the material properties of epithelial tissue have been based on a viscoelastic description. Epithelial tissues have been shown to give rise to both solid and liquid material properties. In addition, numerous cellular properties have been identified that coordinate and regulate cell proliferation. Our vertex model implementation predicts that, as tissues grow, the growth process is dominated by avalanches of cells. However, do epithelial tissues exhibit avalanches *in vivo* under proliferation stress? And which biological and biophysical processes control the onset and magnitude of the avalanches? Addressing these issues is imperative for developing realistic integrated biophysical computational models that accurately describe developmental processes. Especially, the interplay between growth and cellular signaling requires a valid description of the underlying cellular mosaic and its dynamics.

Similar to the signatures identified in our model simulation of avalanche dynamics, time resolved evidence of cell area oscillations have been reported in expanding epithelia (16). However, so far, experiments that involve freely expanding cell cultures have not yet reported this phenomenon. One of the reasons may be the boundary conditions: in freely expanding epithelia, the boundary expands by forming protrusions with cells, often losing confluence. This is in contrast with the simulated system, where the tissue retains its confluency and smooth boundary.

The computational model treats the tissue dynamics within a quasistatic paradigm. That is, it is assumed that the energy relaxation time scale is very short. Changing the solver from gradient descent to overdamped dynamics, will affect the prevalence of avalanches. The magnitude of viscosity, cell properties and the nature of cellular fluctuations control the strength and prevalence of avalanches. When the relaxation time scale increases, the avalanches are smoothed out until they cannot be resolved. *In vitro*, the relaxation time depends on details of the cells, such as cell cycle progression, expression profiles of cadherins and acto-myosin activity. Therefore, the choice of dynamical update rules have profound and fundamental effects on collective phenomena and growth processes, stressing the importance of experimentally quantifying fluctuations and relaxation timescales (63,64). Our computational study illustrates the intricacy of growth processes, their dependence on the dynamics and emergent phenomena when coupling signaling and mechanical responses.

### Computational Methods

#### Spatial Structure

There are many manifestations of vertex models. Here, we adapt a model introduced by Farhadifar et al, which has been successful in emulating the growth of drosophila wing disk epithelia. In this model, the tissue is assumed to evolve quasistatically, i.e., the tissue re-arranges instantaneously to the nearest local minimum via gradient descent (32). This approach originally uses a 2D description of the tissue, but it can be generalized and applied in 3D (36). The energy function considers cell elasticity, actin myosin bundles, and adhesion molecules. Mathematically, it contains an elastic term for area, tensions for each edge and contractility for the perimeter,

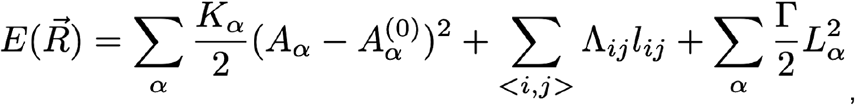

where α is the cell index, *K*_α_ is the elastic coefficient for compressibility, *A*_α_ is the area and *A*_α_^0^ the preferred area, <i,j> is the summation over all edges, Λ _*ij*_ is the edge tension and similarly Γ _α_ is the contractility of the perimeter *L*_α_.

Determining the epithelial tissue structure in a given conformation of cells is more intricate than sole gradient descent energy minimization. Cellular structures undergo topological transitions, if they further minimize the energy of the configuration. The possible transitions have been addressed in the context of foam physics (65). For instance, cells can undergo neighbor exchange via a T1 transition (32,34). Cells do not have a fixed number of edges, and if it is energetically favorable, new edges can form via T3 transitions (32,36). Contrary, cells that shrink undergo inverse T1 transitions, which reduce their number of edges. When a cell obtains three edges or reaches a critical apoptotic size, it can be removed from the structure via a T2 transition (32,65). These transitions conserve the topology of the tissue while implementing energy minimization. The coding environment for the spatial structure was taken from the tyssue repository on github. The tyssue library (DOI 10.5281/zenodo.4475147) is a product of code refactoring originally from (36).

Neglecting the attachment of the eye imaginal disc to the antenna disc and the peripodial membrane (43), we approximate the eye disc to be suspended and with open boundary conditions. Due to the asymmetry of the growth process, as the MF lineary propagates, one cannot implement periodic boundary conditions, as it is often done in other systems (61). To approximate the effect of boundary cells, increased tension was implemented at the cell edges along the boundary. The boundary tension parameters were chosen to enforce an approximately circularity of the final structure SI Fig. 7A. The increased boundary tension did not lead to significant changes in the growth process SI Fig. 7B. This is motivated by the established differential adhesion on tissue boundary layer, often thought as creating an effective surface tension (66,67).

To simulate the structural component of the morphogenetic furrow, a simplistic approach was taken. After determining the cells that make up the MF, their preferred area was instantly set to a decreased value. We term apical constriction factor the percent decrease in preferred cell area of cells in the MF. Furthermore, the preferred area for cells that exit the MF was set back to the default value. In the case of the eye disc model, 2.5 hrs after the cells exhibit the MF, they are artificially made immotile by no longer considering their contribution to the energy term, to simulate the differentiating region.

#### Growth

In the vertex model, cell division can be achieved by directly introducing a new edge in a target cell. Some implementations include area growth, but it has been shown that area does not correlate with volume (29), so we use an alternate implementation. In this implementation, the new edge is chosen by drawing a line through the cell centroid with random orientation (30). The two daughter cells inherit the concentration of the mother cell.

To simulate the cell cycle, a variable V, which corresponds to volume, is assigned to each cell (29). This variable increases over time, and when it crosses a constant threshold, arbitrarily chosen to be 1, the cell divides. To model the mechanical feedback of the cells, also known as mechanosensing, the magnitude of increase over time includes an area dependent term (29). This formulation has been shown to recreate epithelial cell topologies similar to the ones seen in experiments. Finally, to match the experimental cell doubling, the step increase of this is normalized by *t*_*norm*_(*k*), which depends on the growth rate k. The variable and its step increase are shown in the following equation:

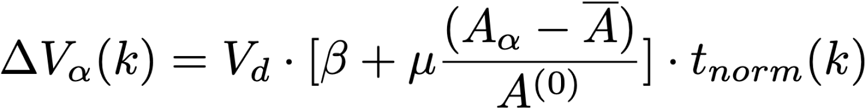

In this equation Δ*V* _α_ is the volume contribution in a single iteration for cell α, *V*_*d*_is the threshold volume for division, β is a constant growth contribution rate, μ determines the strength of mechanosensing contribution to growth and *t*_*norm*_ is a factor that matches the experimental growth rate. Finally *A*_α_, *Ā, A*^(0)^represent the area of cell α, the average cell area in the tissue and the preferred area of the cell respectively. Note that the preferred area of a cell is a structure variable in the vertex model implementation.

The functional form of *t*_*norm*_ depends on the growth profile we want to use. For the exponential growth profile used to model the eye disc, 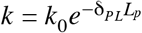 (44,57), where δ _*PL*_ is a constant and *L*_*P*_ is the posterior length (used interchangeably with developmental time since *L*_*P*_ = *v*_*MF*_ *t*). As an approximation, we assume that the target cell has an average area simplifying the equation to:

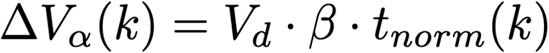

Note that since cells start at *V* _*d*_*/*2 as a starting volume, they require *V _d_/*2 to divide. Now, *t*_*norm*_can be derived from the equation:

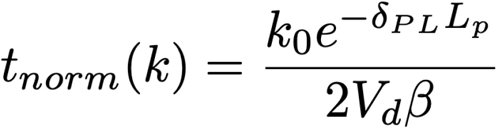

Where in the above, we assume that β and *k*_0_ are in units 1*/s*.

#### Eye Disc Model: Signaling

In order to describe the progression of the MF, a simplified approach was taken. The complete complex interactions of multiple morphogens and transcripts are not needed for this study. To that end, signaling was modeled by considering only a single effective transcription factor obeying an auto-activation regulatory loop. This transcription factor obeys the reaction diffusion equation shown below. The differential equation was discretized on the cell center lattice generated by the vertex model (61). The lattice was approximated by a hexagonal packing and the appropriate discretization method was implemented (68).

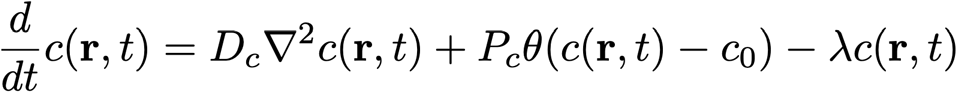

In this equation, c is the concentration of the transcription factor *D*_*c*_ is the diffusion coefficient, *P*_*c*_ is the production rate and λ is the decay rate. The θ depicts a step function which becomes 1 when the concentration crosses *c*_0_. The emergent dynamics of this equation, after initializing a concentration or source, give rise to a non-dissipating linearly propagating concentration gradient. This is ideal for the eye disc, as the MF is known to propagate at a constant speed of 3.4μ*m/hr*, as measured by the dorsal-ventral point of the MF (44). Finally, cells that constitute the MF were defined as cells that have a concentration between a range, chosen to be 0.4 and 1.

Given the autoregulatory loop for MF propagation, an initial flux or concentration of the effective morphogen is necessary. *In vitro*, Hh diffuses through the posterior tissue boundary and initiates the signaling cascade necessary for MF propagation. To initialize the propagation of the MF in the eye disc simulation, we follow a similar approach with (57). The posterior boundary cells obeying *x*(*t*) ≤ 0.2 *L*_*AP*_, where *L*_*AP*_ is the total anterior to posterior length, received a boundary flux of the morphogen equal to 40% of the autoproduction rate *P*_*c*_. Varying this parameter simply changes the time scale of the initialization phase. Furthermore, the boundary flux was tapered down by a time dependent term τ(*t*) = 1 – θ(*t* – 10*hr*), such that the boundary flux ceases after 10 hrs.

#### Model calibration and initialization

The specifications to initialize the eye disc size and shape were taken from (57). An elliptical tissue was extracted from a previously grown epithelial tissue under periodic boundary conditions as in (29,32). For the uniform growth model, a circular tissue with a radius of 11μm was extracted from the same previously grown epithelial tissue. At the start of both uniform growth and eye disc simulations, cells are set to be at a random point of their cell cycle. That is done by setting the variable V, defined above, at a random value from 0.5 to 1. Note when V=1, a cell divides and V is halved, making the minimum V be 0.5.

Next, we discuss how we set the spatial and temporal scales in the simulation. Firstly, the time and space scales used in the generic uniform growth model were taken from the eye disc. Given that the average cell size in the posterior portion of the eye disc is known to be 9μ*m*^2^ (47), obtaining the average cell area from a sample simulation and equating it to the experimental value provides the length scale. For an intermediate value of constriction, the MF propagation speed was measured. Then, it was equated to the experimentally derived value of 3.4 μ*m /hr* (44) to obtain the time scale. This allowed for the conversion of simulation time to developmental time and thus, the calibration of the growth rate. The apical constriction is known to cause the 9 μ*m*^2^cells ahead of the furrow to 0.6 μ*m*^2^(47). The two dimensional vertex model cannot support such an intense constriction. To this end the apical constriction is varied and the corresponding emergent behaviors are investigated.

#### Coupling signaling and tissue structure iterations

A custom script was developed for the eye disc model, which combined reaction diffusion based signaling and the vertex model. Firstly, the tyssue library was used for the vertex model and its dynamical updates. For signaling, which translates to ordinary differential equations under discretization, python’s scipy.integrate.solve_ivp() function was used. The two aspects were updated sequentially. The differential equations were time-evolved for a constant interval of time. Then, the properties of the cells, a subset of which depend on the morphogen concentration, were updated. Afterwards, the tyssue solver was used to relax the system to the nearest energy minimum. Then, the configuration of the resulting structure was plugged in the differential equations and they were time propagated. The time resolution of this transition between signaling evolution and mechanical update was increased until the macroscopic system observables were invariant. The mechanical structure and cell state were updated every 4 minutes of developmental time.

## Supplemental information

### Supplemental Movies

SI Vid 1 [winglike_growth.avi](https://drive.google.com/file/d/1fDACO-5u8V220lvyaYtKv9FJXHt3Ssmo/view)

SI Vid 2 [eye_growth.avi](https://drive.google.com/file/d/1QdHidFOGKxDI9NuvnjEzZ56xcnErCJ0F/view)

## Supplemental Figures

**SI Fig. 1.**
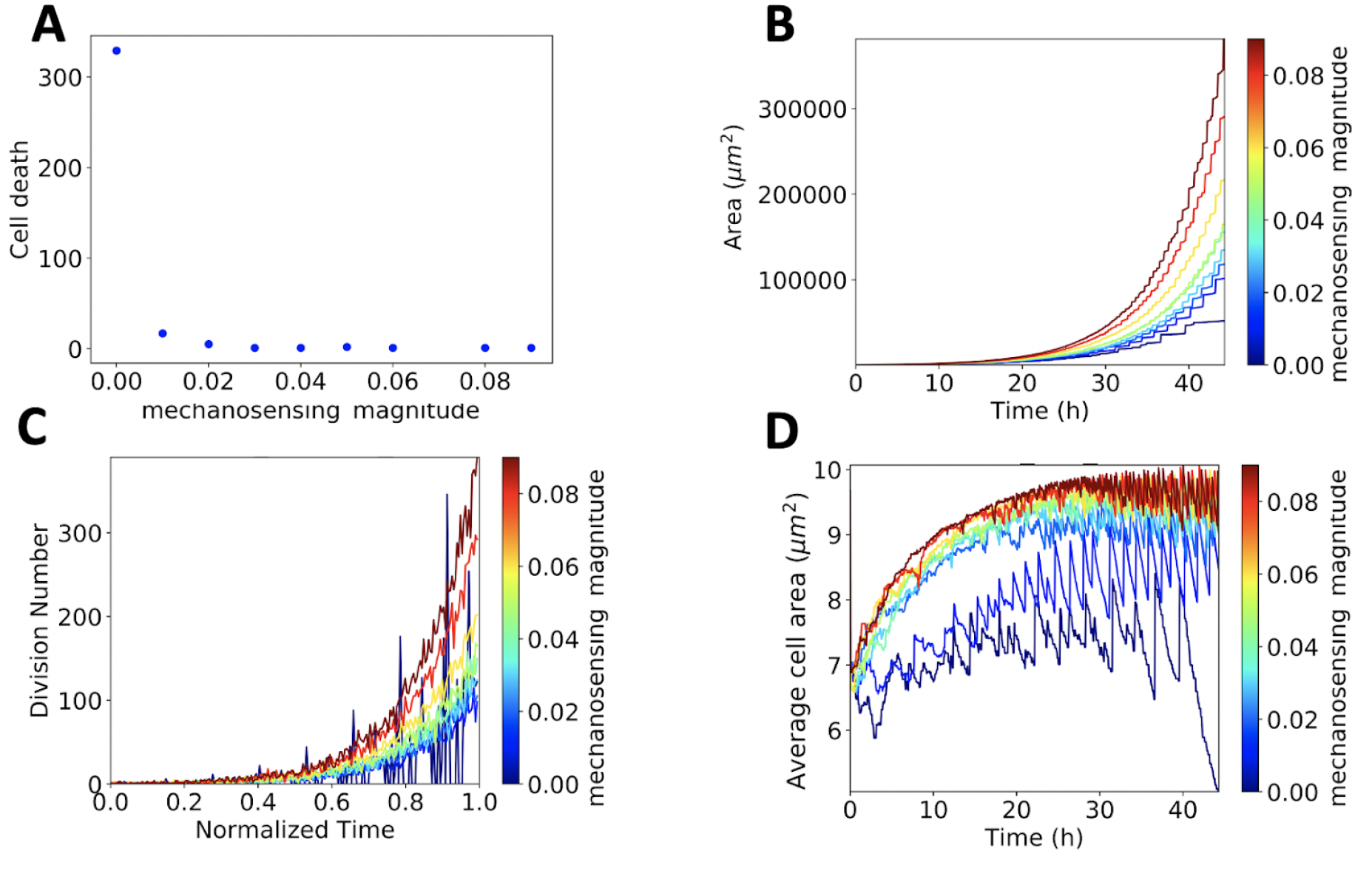
Mechanosensing analysis in the uniform growth model. **A:** Cell death during growth for different mechanosensing amplitudes. When mechanosensing is toggled off, at zero mechanosensing magnitude (μ), there is significant cell death due to proliferation stresses. When non-zero values of mechanosensing are used, cell death becomes negligible. **B:** Area versus time for different mechanosensing amplitudes. The mechanosensing amplitude significantly affects the growth process. Since this is exponential growth, small variations intensify exponentially. Close to μ=0.04, which is the parameter used and is taken from (29), the area does not deviate when changing mechanosensing. Growth discontinuities or avalanches are present for all mechanosensing parameters. **C)** Division number versus time for variable mechanosensing magnitudes. When mechanosensing is turned off, the cells divide periodically since all daughter cells grow instantaneously. As the mechanosensing amplitude increases, the division number increases. That is the case because ΔV is bounded by 0, meaning cells can only grow or arrest their cell cycle, making the ΔV contribution due to mechanosensing biased to be positive. **D)** Average cell area versus time for different mechanosensing amplitudes. When mechanosensing is off, the tissue does not reach the preferred area of 9 μ*m*^2^.

**SI Fig. 2.**
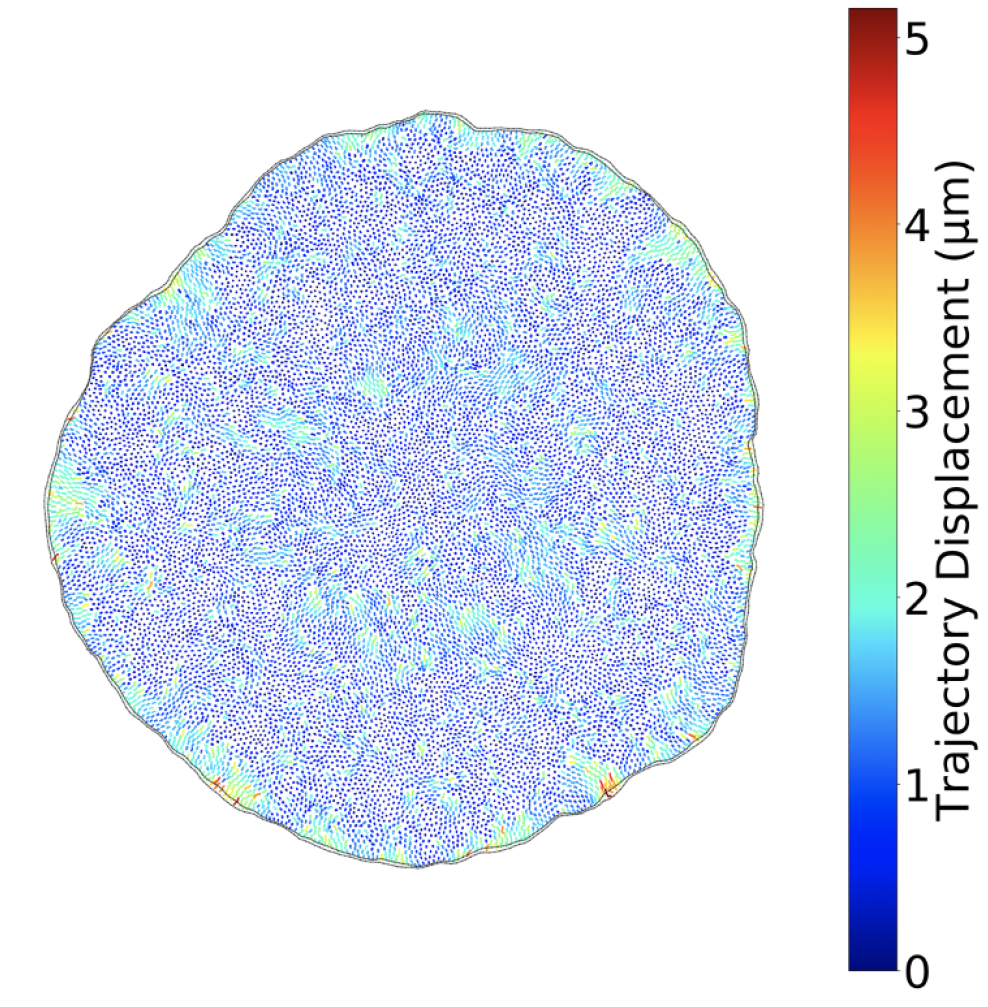
Cell trajectories between two successive avalanches. The movement of cells recorded after an avalanche at T=44.1h and before the next avalanche at T=44.9h. Cell movement does not exhibit the collective behavior seen during an avalanche. There are disparate regions of the tissue where cells exhibit significant displacement and the boundary does not significantly expand.

**SI Fig. 3.**
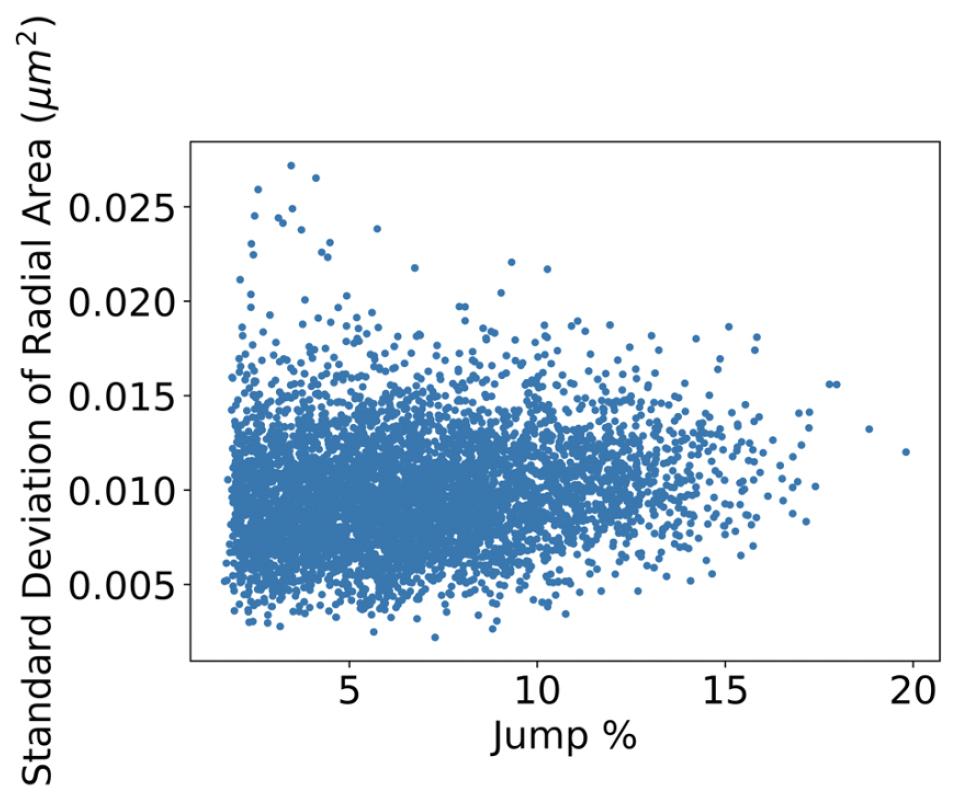
Standard deviation of the radial average cell area right before an avalanche versus the tissue jump magnitude after the onset of the given avalanche. The y-axis provides a measure of cell packing inhomogeneity. The correlation was found to be weak but non-negligible, with a spearman’s rank correlation coefficient of ρ=0.17 and a p-value of 0.

**SI Fig. 4.**
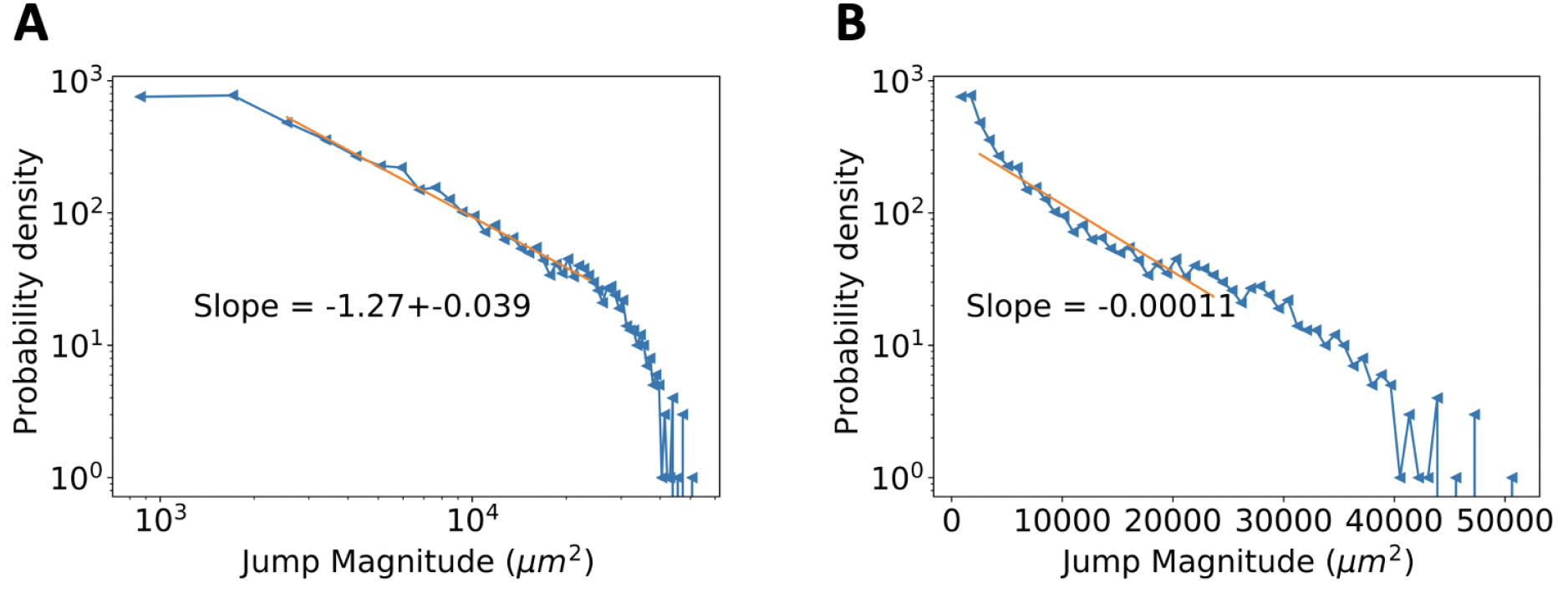
Fitting considerations. **A:** Log-log plot of the probability density versus tissue jump (or avalanche) magnitude. The coefficient was obtained to be τ=1.27 by a linear fit. The power law has an exponential cutoff due to finite system size. **B:** Semi-log plot for probability density versus tissue jump magnitude. The linear fit is poor as the distribution does not exhibit an exponential signature.

**SI Fig. 5.**
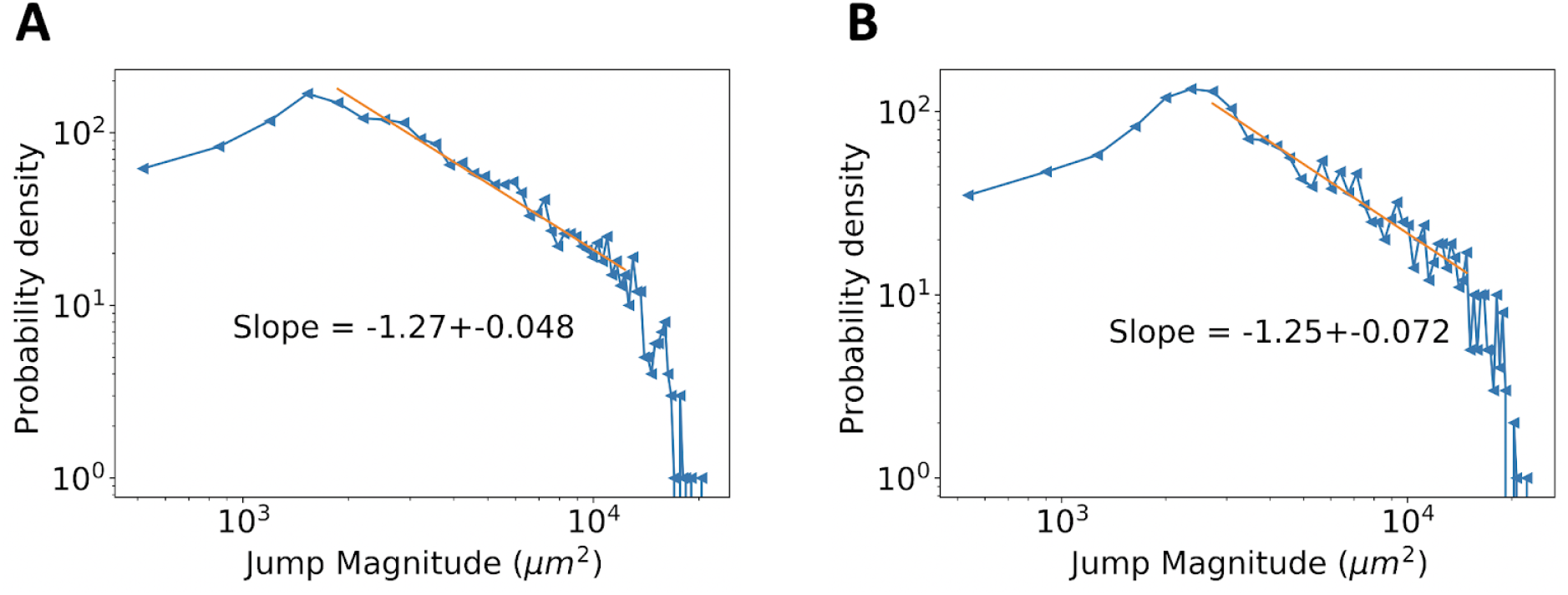
Exponent dependence on parameter variation. **A:** Log-log plot of the probability density versus tissue jump magnitude when mechanosensing amplitude, μ, is one quarter of the original value. The slope coefficient was obtained to be invariant with τ=1.27. **B:** Log-log plot of the probability density versus tissue jump magnitude when boundary tension is changed to the bulk edge tension. The slope coefficient was obtained to be τ=1.25 and considering the error bar, within the 1.27 value.

**SI Fig. 6.**
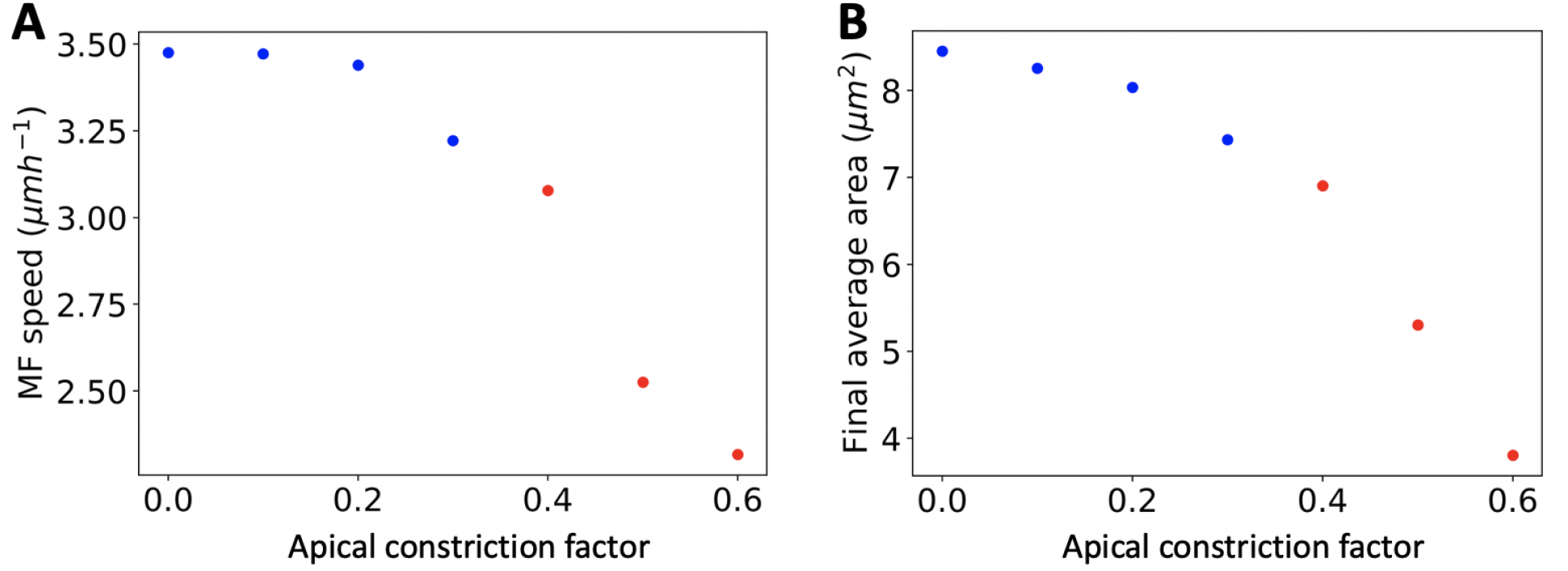
Effect of apical constriction on MF speed and cell packing. **A:** The MF speed decreases as apical constriction increases. **B:** The cellular mosaic becomes increasingly packed with increasing apical constriction. This speed dependence on apical constriction is a consequence of the signaling differential equations, which are discretized on the cellular lattice, and are independent of the physical distance between cells. The blue dots involve data taken from simulations were the MF reached the anterior portion and the growth process was concluded. The red dots indicate data from simulations that were not completed since the tissue reaches a very large number of cells, making them very computationally costly.

**SI Fig. 7.**
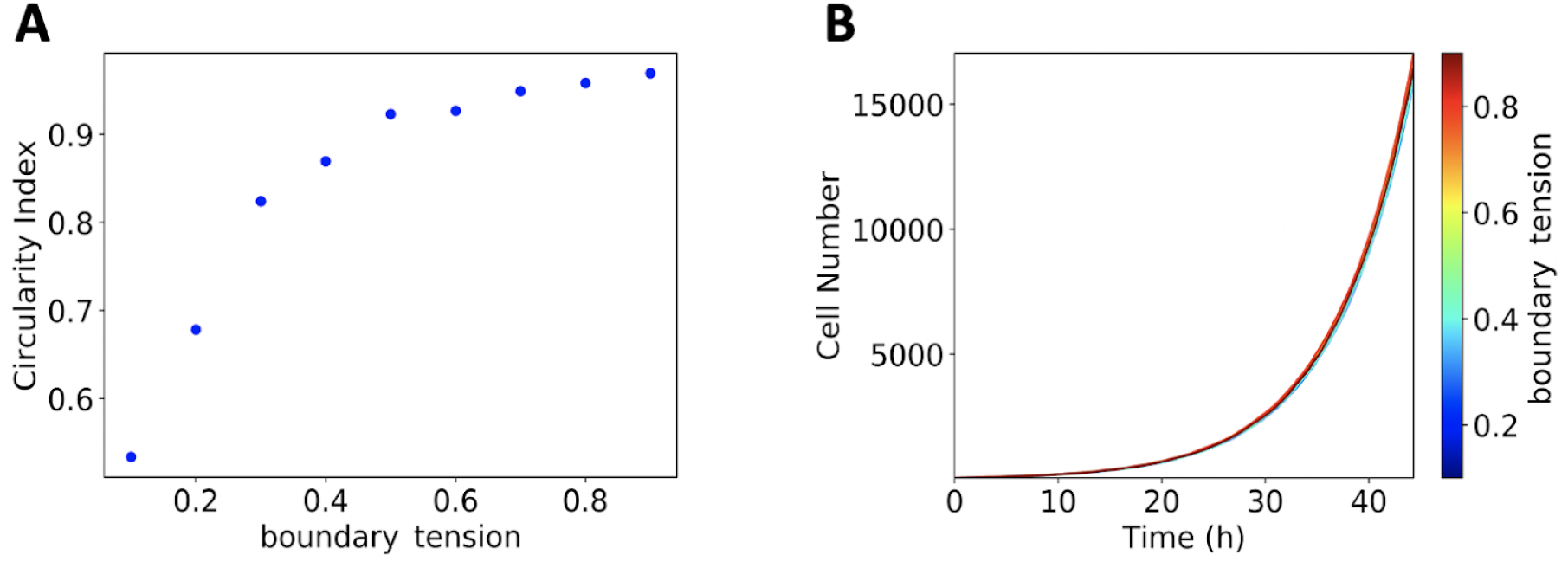
Boundary tension analysis. **A:** Circularity index, defined as *c*_*0*_ = 4π*AP* ^2^, where A is the tissue area and P the perimeter. The tissue became increasingly circular as the boundary tension was increased. The effect of increasing boundary tension is more prominent for low values and saturates ∼1 for larger values. Boundary tension value used throughout this paper was set to 0.7. **B:** Cell number versus time. The changes in boundary tension do not significantly affect the growth rate of the tissue.

## Acknowledgements

We would like to thank Guillaume Gay, Lisa Manning, Yehuda Ben-Zion, Konstantin Kozlov, Svetlana Surkova, and Leonardo Morsut for useful discussions. Research reported in this publication was supported by the National Institutes of Health under award number NIH GM128193. The content is solely the responsibility of the authors and does not necessarily represent the official views of the National Institutes of Health. Computation for the work described in this paper was supported by the University of Southern California’s Center for High-Performance Computing (carc.usc.edu).

## Notes

### Competing Interest Statement

The authors have declared no competing interest.

### Summary of Updates

Cellular vortex formation

## References

1. Forgacs G, Foty RA, Shafrir Y, Steinberg MS. Viscoelastic Properties of Living Embryonic Tissues: a Quantitative Study. Biophys J. 1998 May 1;74(5):2227–34.

2. Schötz E-M, Lanio M, Talbot JA, Manning ML. Glassy dynamics in three-dimensional embryonic tissues. J R Soc Interface. 2013 Dec 6;10(89):20130726.

3. Angelini TE, Hannezo E, Trepat X, Marquez M, Fredberg JJ, Weitz DA. Glass-like dynamics of collective cell migration. Proc Natl Acad Sci. 2011 Mar 22;108(12):4714–9.

4. Henkes S, Fily Y, Marchetti MC. Active jamming: Self-propelled soft particles at high density. Phys Rev E. 2011 Oct 12;84(4):040301.

5. Berthier L. Nonequilibrium Glassy Dynamics of Self-Propelled Hard Disks. Phys Rev Lett. 2014 Jun 4;112(22):220602.

6. Berthier L, Kurchan J. Non-equilibrium glass transitions in driven and active matter. Nat Phys. 2013 May;9(5):310–4.

7. Bi D, Lopez JH, Schwarz JM, Manning ML. A density-independent glass transition in biological tissues. Nat Phys. 2015 Dec;11(12):1074–9.

8. Ruina A. Slip instability and state variable friction laws. J Geophys Res Solid Earth. 1983;88(B12):10359–70.

9. Hubert A, Schäfer R. Magnetic Domains: The Analysis of Magnetic Microstructures [Internet]. Berlin Heidelberg: Springer-Verlag; 1998 [cited 2021 Jan 8]. Available from: https://www.springer.com/gp/book/9783540641087

10. Bak P, Tang C, Wiesenfeld K. Self-organized criticality: An explanation of the 1/ f noise. Phys Rev Lett. 1987 Jul 27;59(4):381–4.

11. Karimi K, Ferrero EE, Barrat J-L. Inertia and universality of avalanche statistics: The case of slowly deformed amorphous solids. Phys Rev E. 2017 Jan 12;95(1):013003.

12. Salerno KM, Maloney CE, Robbins MO. Avalanches in Strained Amorphous Solids: Does Inertia Destroy Critical Behavior? Phys Rev Lett. 2012 Sep 6;109(10):105703.

13. Bohn F, Durin G, Correa MA, Machado NR, Pace RDD, Chesman C, et al. Playing with universality classes of Barkhausen avalanches. Sci Rep. 2018 Jul 26;8(1):1–12.

14. Popovic M, Druelle V, Dye N, Jülicher F, Wyart M. Inferring the flow properties of epithelial tissues from their geometry. New J Phys [Internet]. 2020 [cited 2021 Feb 17]; Available from: http://iopscience.iop.org/article/10.1088/1367-2630/abcbc7

15. Streichan SJ, Hoerner CR, Schneidt T, Holzer D, Hufnagel L. Spatial constraints control cell proliferation in tissues. Proc Natl Acad Sci. 2014 Apr 15;111(15):5586–91.

16. Serra-Picamal X, Conte V, Vincent R, Anon E, Tambe DT, Bazellieres E, et al. Mechanical waves during tissue expansion. Nat Phys. 2012 Aug;8(8):628–34.

17. Vedula SRK, Leong MC, Lai TL, Hersen P, Kabla AJ, Lim CT, et al. Emerging modes of collective cell migration induced by geometrical constraints. Proc Natl Acad Sci. 2012 Aug 7;109(32):12974–9.

18. Doxzen K, Vedula SRK, Leong MC, Hirata H, Gov NS, Kabla AJ, et al. Guidance of collective cell migration by substrate geometry. Integr Biol. 2013 Aug 22;5(8):1026–35.

19. Puliafito A, Hufnagel L, Neveu P, Streichan S, Sigal A, Fygenson DK, et al. Collective and single cell behavior in epithelial contact inhibition. Proc Natl Acad Sci. 2012 Jan 17;109(3):739–44.

20. Heinrich MA, Alert R, LaChance JM, Zajdel TJ, Košmrlj A, Cohen DJ. Size-dependent patterns of cell proliferation and migration in freely-expanding epithelia [Internet]. eLife. eLife Sciences Publications Limited; 2020 [cited 2021 Jan 15]. Available from: https://elifesciences.org/articles/58945

21. Huergo MAC, Pasquale MA, González PH, Bolzán AE, Arvia AJ. Dynamics and morphology characteristics of cell colonies with radially spreading growth fronts. Phys Rev E. 2011 Aug 11;84(2):021917.

22. Lee RM, Kelley DH, Nordstrom KN, Ouellette NT, Losert W. Quantifying stretching and rearrangement in epithelial sheet migration. New J Phys. 2013 Feb;15(2):025036.

23. Eder D, Aegerter C, Basler K. Forces controlling organ growth and size. Mech Dev. 2017 Apr 1;144:53–61.

24. Pan Y, Heemskerk I, Ibar C, Shraiman BI, Irvine KD. Differential growth triggers mechanical feedback that elevates Hippo signaling. Proc Natl Acad Sci. 2016 Nov 8;113(45):E6974–83.

25. Irvine KD, Shraiman BI. Mechanical control of growth: ideas, facts and challenges. Development. 2017 Dec 1;144(23):4238–48.

26. Ehrig S, Schamberger B, Bidan CM, West A, Jacobi C, Lam K, et al. Surface tension determines tissue shape and growth kinetics. Sci Adv. 2019 Sep 1;5(9):eaav9394.

27. Mao Y, Tournier AL, Hoppe A, Kester L, Thompson BJ, Tapon N. Differential proliferation rates generate patterns of mechanical tension that orient tissue growth. EMBO J. 2013 Oct 30;32(21):2790–803.

28. Gibson WT, Veldhuis JH, Rubinstein B, Cartwright HN, Perrimon N, Brodland GW, et al. Control of the Mitotic Cleavage Plane by Local Epithelial Topology. Cell. 2011 Feb 4;144(3):427–38.

29. Aegerter-Wilmsen T, Smith AC, Christen AJ, Aegerter CM, Hafen E, Basler K. Exploring the effects of mechanical feedback on epithelial topology. Development. 2010 Feb 1;137(3):499–506.

30. Aegerter-Wilmsen T, Heimlicher MB, Smith AC, de Reuille PB, Smith RS, Aegerter CM, et al. Integrating force-sensing and signaling pathways in a model for the regulation of wing imaginal disc size. Development. 2012 Sep 1;139(17):3221–31.

31. Salbreux G, Barthel LK, Raymond PA, Lubensky DK. Coupling Mechanical Deformations and Planar Cell Polarity to Create Regular Patterns in the Zebrafish Retina. PLOS Comput Biol. 2012 Aug 23;8(8):e1002618.

32. Farhadifar R, Röper J-C, Aigouy B, Eaton S, Jülicher F. The Influence of Cell Mechanics, Cell-Cell Interactions, and Proliferation on Epithelial Packing. Curr Biol. 2007 Dec;17(24):2095–104.

33. Alt Silvanus, Ganguly Poulami, Salbreux Guillaume. Vertex models: from cell mechanics to tissue morphogenesis. Philos Trans R Soc B Biol Sci. 2017 May 19;372(1720):20150520.

34. Fletcher AG, Osterfield M, Baker RE, Shvartsman SY. Vertex Models of Epithelial Morphogenesis. Biophys J. 2014 Jun;106(11):2291–304.

35. Barton DL, Henkes S, Weijer CJ, Sknepnek R. Active Vertex Model for cell-resolution description of epithelial tissue mechanics. PLOS Comput Biol. 2017 Jun 30;13(6):e1005569.

36. Monier B, Gettings M, Gay G, Mangeat T, Schott S, Guarner A, et al. Apico-basal forces exerted by apoptotic cells drive epithelium folding. Nature. 2015 Feb;518(7538):245–8.

37. Lewis FT. The correlation between cell division and the shapes and sizes of prismatic cells in the epidermis of cucumis. Anat Rec. 1928;38(3):341–76.

38. Heller D, Hoppe A, Restrepo S, Gatti L, Tournier AL, Tapon N, et al. EpiTools: An Open-Source Image Analysis Toolkit for Quantifying Epithelial Growth Dynamics. Dev Cell. 2016 Jan 11;36(1):103–16.

39. Sánchez-Gutiérrez D, Tozluoglu M, Barry JD, Pascual A, Mao Y, Escudero LM. Fundamental physical cellular constraints drive self-organization of tissues. EMBO J. 2016 Jan 4;35(1):77–88.

40. Escudero LM, Costa L da F, Kicheva A, Briscoe J, Freeman M, Babu MM. Epithelial organisation revealed by a network of cellular contacts. Nat Commun. 2011 Nov 8;2(1):1–7.

41. Patel AB, Gibson WT, Gibson MC, Nagpal R. Modeling and Inferring Cleavage Patterns in Proliferating Epithelia. PLOS Comput Biol. 2009 Jun 12;5(6):e1000412.

42. Kumar JP. My what big eyes you have: How the Drosophila retina grows. Dev Neurobiol. 2011 Dec;71(12):1133–52.

43. Kumar JP. The fly eye: Through the looking glass. Dev Dyn. 2018;247(1):111–23.

44. Vollmer J, Fried P, Sánchez-Aragón M, Lopes CS, Casares F, Iber D. A quantitative analysis of growth control in the Drosophila eye disc. Development. 2016 May 1;143(9):1482–90.

45. Kumar JP. Building an ommatidium one cell at a time. Dev Dyn. 2012;241(1):136–49.

46. Treisman JE. Retinal differentiation in Drosophila. Wiley Interdiscip Rev Dev Biol. 2013;2(4):545–57.

47. Wolff T, Ready DF. The beginning of pattern formation in the Drosophila compound eye: the morphogenetic furrow and the second mitotic wave. Development. 1991 Nov 1;113(3):841–50.

48. Baker NE. Cell proliferation, survival, and death in the Drosophila eye. Semin Cell Dev Biol. 2001 Dec;12(6):499–507.

49. Crossman SH, Streichan SJ, Vincent J-P. EGFR signaling coordinates patterning with cell survival during Drosophila epidermal development. PLOS Biol. 2018 Oct 31;16(10):e3000027.

50. Vollmer J, Fried P, Aguilar-Hidalgo D, Sánchez-Aragón M, Iannini A, Casares F, et al. Growth control in the https://www.zotero.org/google-docs/?7TMgbN eye disc by the cytokine Unpaired. Development. 2017 Mar 1;144(5):837–43.

51. Curtiss J, Mlodzik M. Morphogenetic furrow initiation and progression during eye development in Drosophila: the roles of decapentaplegic, hedgehog and eyes absent. Development. 2000;127(6):1325–36.

52. Domínguez M, Hafen E. Hedgehog directly controls initiation and propagation of retinal differentiation in the Drosophila eye. Genes Dev. 1997 Dec 1;11(23):3254–64.

53. Heberlein U, Moses K. Mechanisms of Drosophila retinal morphogenesis: the virtues of being progressive. Cell. 1995 Jun 30;81(7):987–90.

54. Pichaud F, Casares F. homothorax and iroquois-C genes are required for the establishment of territories within the developing eye disc. Mech Dev. 2000 Aug;96(1):15–25.

55. Roignant J-Y, Treisman JE. Pattern formation in the Drosophila eye disc. Int J Dev Biol. 2009 May 22;53(5–6):795–804.

56. Brown NL, Sattler CA, Paddock SW, Carroll SB. Hairy and Emc negatively regulate morphogenetic furrow progression in the drosophila eye. Cell. 1995 Mar 24;80(6):879–87.

57. Fried P, Sánchez-Aragón M, Aguilar-Hidalgo D, Lehtinen B, Casares F, Iber D. A Model of the Spatio-temporal Dynamics of Drosophila Eye Disc Development. Thieffry D, editor. PLOS Comput Biol. 2016 Sep 14;12(9):e1005052.

58. Okuda S, Miura T, Inoue Y, Adachi T, Eiraku M. Combining Turing and 3D vertex models reproduces autonomous multicellular morphogenesis with undulation, tubulation, and branching. Sci Rep. 2018 Feb 5;8(1):2386.

59. Schilling S, Willecke M, Aegerter-Wilmsen T, Cirpka OA, Basler K, Mering C von. Cell-Sorting at the A/P Boundary in the Drosophila Wing Primordium: A Computational Model to Consolidate Observed Non-Local Effects of Hh Signaling. PLOS Comput Biol. 2011 Apr 7;7(4):e1002025.

60. Smith AM, Baker RE, Kay D, Maini PK. Incorporating chemical signalling factors into cell-based models of growing epithelial tissues. J Math Biol Heidelb. 2012 Sep;65(3):441–63.

61. Wartlick O, Mumcu P, Kicheva A, Bittig T, Seum C, Jülicher F, et al. Dynamics of Dpp Signaling and Proliferation Control. Science. 2011 Mar 4;331(6021):1154–9.

62. Talamali M, Petäjä V, Vandembroucq D, Roux S. Avalanches, precursors, and finite-size fluctuations in a mesoscopic model of amorphous plasticity. Phys Rev E. 2011 Jul 29;84(1):016115.

63. Fodor É, Mehandia V, Comelles J, Thiagarajan R, Gov NS, Visco P, et al. Spatial Fluctuations at Vertices of Epithelial Layers: Quantification of Regulation by Rho Pathway. Biophys J. 2018 Feb 27;114(4):939–46.

64. Curran S, Strandkvist C, Bathmann J, de Gennes M, Kabla A, Salbreux G, et al. Myosin II Controls Junction Fluctuations to Guide Epithelial Tissue Ordering. Dev Cell. 2017 Nov 20;43(4):480-492.e6.

65. Stavans J. The evolution of cellular structures. Rep Prog Phys. 1993 Jun 1;56(6):733–89.

66. Sussman DM, Schwarz JM, Marchetti MC, Manning ML. Soft yet Sharp Interfaces in a Vertex Model of Confluent Tissue. Phys Rev Lett. 2018 Jan 29;120(5):058001.

67. Dahmann C, Oates AC, Brand M. Boundary formation and maintenance in tissue development. Nat Rev Genet. 2011 Jan;12(1):43–55.

68. Fabero JC, Bautista A, Casasús L. An explicit finite differences scheme over hexagonal tessellation. Appl Math Lett. 2001 Jan 1;14(5):593–8.

